# Bipartite DNA binding domain of transcription factor BCL11B binds clustered short DNA sequence motifs

**DOI:** 10.64898/2026.05.01.721897

**Authors:** Jisun Lee, Jujun Zhou, John R. Horton, Meigen Yu, Munachi Daniella Muoghalu, Faizan Ahmed Khan, Xiaotian Zhang, Yun Huang, Robert M. Blumenthal, Xing Zhang, Xiaodong Cheng

## Abstract

<u>B</u>-<u>c</u>ell <u>l</u>eukemia/lymphoma 11B (BCL11B), despite its name, is a key regulator of T-cell development, specification, and T-cell malignancies. BCL11B contains a bipartite DNA binding domain composed of two C2H2 zinc finger arrays: low-affinity ZF2-3 and high affinity ZF4-6. These arrays function as homotypic modules that recognize similar six-nucleotide motifs, TG(n)CC(c/t/a), as seven of the eight DNA base-contacting residues are conserved between them. The most conserved interactions involve GG dinucleotides, contacted by arginine and lysine residues at key base-interacting positions in ZF3 and ZF5. The two ZF arrays are connected by a long ∼300-residue linker that provides flexibility in how the arrays engage DNA, allowing ZF2-3 and ZF4-6 binding to the same or opposite strands with variable orientation, spacing and positioning along the DNA. This extended linker is enriched in serine/threonine, acidic residues (aspartate/glutamate), and structural residues (glycine/proline), providing additional layers of transcriptional regulation possibly through post-translational modification, electrostatic modulation, and/or condensate formation. We also examined six missense mutations in base-interacting residues, that are associated with neurodevelopmental disorders. Substitutions replacing bulky, positively charged arginine or lysine with smaller or hydrophobic residues likely reduce DNA-binding affinity and/or specificity, whereas substitutions between asparagine and lysine may alter base recognition preferences.

## Introduction

<u>B</u>-<u>c</u>ell <u>l</u>eukemia/lymphoma 11A (BCL11A) and 11B (BCl11B) are paralogous DNA-binding transcription factors, that each contain two arrays of C2H2 zinc fingers (ZF2-ZF3 and ZF4-ZF6). BCL11<u>A</u> is vital for B-cell development ^1,2^ and for the postnatal switch from fetal-to-adult hemoglobin in erythroid cells ^3^. Despite its misleading name, BCL11<u>B</u> is neither expressed in B-cell chronic lymphocytic leukemia nor in B-cell lymphoma ^4,5^. Instead, BCL11B is crucial for T-cell development and specification ^6–12^, and plays a role in T-cell malignancies ^13–15^.

These two factors also jointly regulate the production and differentiation of cortical projection neurons in mice, with redundant and compensatory expression ^16^. Bcl11a is predominantly expressed in Purkinje cells of both the developing and adult mouse cerebellum, and Bcl11a deficiency in cerebellar Purkinje cells causes ataxia and autistic-like behavior ^17^. Loss-of-function experiments in *Bcl11b* null mutant mice demonstrate that Bcl11b plays a critical role in the development of corticospinal motor neurons axonal projections to the spinal cord *in vivo* ^18^. A tetracycline-dependent inducible mouse model revealed that adult expression of Bcl11b is essential for the survival, differentiation, and functional integration of adult-born granule neurons, and is also required for the survival of pre-existing mature neurons ^19^. Consequently, selective loss of Bcl11b expression in the adult hippocampus results in impaired spatial working memory ^19^. In a mouse striatal precursor cell line, Bcl11b target genes are significantly associated with brain-derived neurotrophic factor signaling ^20^.

Furthermore, mutations in these genes have been linked to neurodevelopmental disorders in humans. In a cohort study of 77 individuals with heterozygous variants in BCL11<u>A</u>, 60 unique disease-associated variants were identified, including frameshift, stop-gain/nonsense, microdeletions, missense, and splice-site mutations ^21^. These variants are distributed across the entire length of the protein. Four missense variants are located in the N-terminal region shared by the three major isoforms (-S, -L, and -XL), whereas two C-terminal missense variants fall within the DNA binding zinc fingers ZF4 (N746K) and ZF5 (K784T) unique to the largest -XL isoform. Because BCL11A plays a key role in silencing fetal hemoglobin, the most common clinical features associated with BCL11A mutations include persistence of fetal hemoglobin and hindbrain abnormalities ^21^. Notably, BCL11A haploinsufficiency is associated with neurodevelopmental disorders, and is also known as Dias-Logan syndrome ^22^. A related condition, 2p15-p16.1 microdeletion syndrome ^23^, involves chromosomal deletions that can affect multiple genes, including BCL11A ^24,25^.

In a parallel study of 92 individuals carrying BCL11<u>B</u> germline variants, 39 (51 total mutants and 12 duplications) unique variants were identified including 16 missense mutations ^26^. Significantly, all identified missense variants were located within the DNA binding zinc fingers: one in ZF2, two in ZF3, three in ZF4, nine in ZF5, and one in ZF6. A representative case is a newborn boy with a missense mutation, Asn441-to-Lys (N441K) in ZF2 of BCL11B, who presented with severe combined immunodeficiency characterized by T-lymphocyte deficiency but normal B and natural killer lymphocytes ^27^. [In T^-^B^+^ severe combined immunodeficiency (SCID), B cells are present but do not develop properly in the absence of T-cell help ^28^.] Genotype-phenotype analyses suggest that phenotypic severity and variability largely depend on the DNA-binding affinity and specificity of the altered BCL11B proteins ^26^. Furthermore, a distinct DNA methylation pattern associated with BCL11B mutant variants may serve as a diagnostic biomarker ^29^. In cancer, dysregulation of *BCL11B* can also arise through structural rearrangement. In a subset of acute leukemias, chromosomal rearrangements place *BCL11B* under the control of distal super-enhancers, leading to aberrant allele-specific expression ^30^. Conversely, in t(5;14) leukemia, a distal *BCL11B* enhancer is hijacked to activate the homeobox TLX3 oncogene ^31^.

Given the unusual importance of these two transcription factors in hematological, immunological, and neurological development, in this study we compare the DNA binding properties of BCL11A and BCL11B. They share a highly conserved overall domain organization, as well as strong sequence conservation within their DNA-binding zinc finger arrays. Both proteins contain two ZF tandem arrays (ZF2-3 and ZF4-6) separated by an approximately 300-residue spacer (Figure 1A). Previous structural investigations have primarily focused on the higher-affinity binding of the ZF4-6 array of BCL11A to the DNA motif TGnCCA (n = any nucleotide) derived from the globin locus ^32–34^, as well as the low-affinity interaction between the ZF2-3 array of BCL11A and the related sequence TGtCCA ^35^. Here, we investigate the structural basis for DNA binding specificity of BCL11B, with particular emphasis on the ZF2-ZF3 array. Upon examining published ENCODE ChIP-seq data of eGFP-BCL11B in human HEK293 cells ^36^, we observed that multiple copies of these homotypic motifs are frequently present within BCL11B binding sites. We therefore propose that cooperative DNA binding by the ZF tandem arrays of homodimers, or BCL11B-BCL11A heterodimers, may extend the length of the recognized DNA sequence, thereby enhancing the specificity and/or sensitivity of transcriptional regulation. We test this hypothesis via structural and DNA-protein interaction analyses.

**Figure 1.**
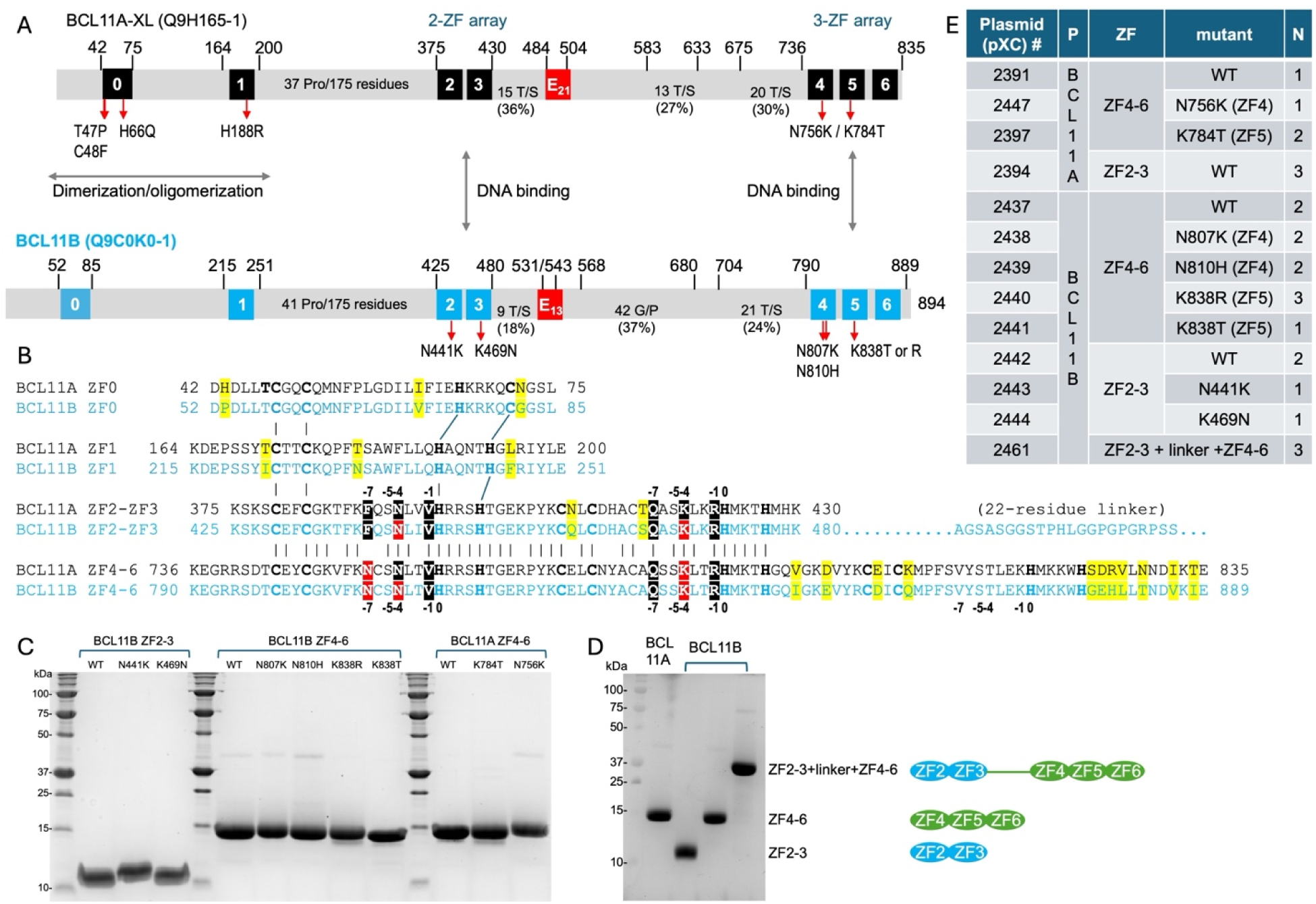
BCL11A and BCL11B share a similar overall domain organization. (**A**) Schematic representations of the BCL11A-XL isoform and full-length BCL11B, with corresponding UniProt accession numbers indicated. Individual C2H2-ZF units are numbered (ZF0-ZF6) and highlighted in black or cyan. Acidic regions are labeled E in red boxes, and proline-rich, glycine/proline-rich, and serine/threonine-rich regions are indicated. Selected missense mutations in BCL11A and BCL11B, associated with neurodevelopmental disorders, are indicated within ZF units. (**B**) Pairwise sequence alignments of BCL11A and BCL11B across ZF0, ZF1, ZF2-3, and ZF4-6. For each ZF unit, DNA base-interacting positions are denoted as -1, -4, -5 and -7 relative to the first zinc-coordinating histidine (position 0). Residues associated with pathogenic mutations in ZF2-3 and ZF4-5 are highlighted in red. (**C**) Examples of recombinant proteins of BCL11B ZF2-3 and ZF4-6 (WT and mutants) used in this study. (**D**) Examples of recombinant BCL11B proteins of isolated ZF2-3, ZF4-6, and a linked ZF2-3 and ZF4-6 construct connected by a 22-residue linker, containing seven glycine, four proline and five serine residues. (**E**) Summary of expression constructs for the ZF arrays used in this study. N is the number of independent purifications.

## Results

### Comparison between BCL11A and BCL11B

Human BCL11A and BCL11B each exist in multiple isoforms (UniProt entries Q9H165 and Q9C0K0) that differ in the number of C-terminal zinc fingers ^21,34,37–39^. Here we focus on the largest isoforms, BCL11A-XL (835 residues) and full-length BCL11B (894 residues) (Figure 1A). Although these two proteins share a similar overall domain organization, they exhibit just over 60% (509/835 residues) sequence identity (Figure S1A). The invariant residues are concentrated within the ZFs: the N-terminal single fingers ZF0 (91% = 30/34 residues) and ZF1 (92% = 34/37), as well as the tandem arrays ZF2-3 (96% = 54/56) and ZF4-6 (89% = 89/100) (Figure 1B).

The respective N-terminal regions of BCL11A ^34,40^ and BCL11B ^41–43^ have been implicated in oligomerization. While the isolated ZF0 of BCL11A forms a tetramer in the crystal structure ^40^, solution NMR studies indicate that ZF0 exists predominantly as a monomer even at 40 μM concentrations ^44^. Oligomerization has been proposed as a mechanism underlying the effects of heterozygous missense mutations, wherein mutant proteins can act in a dominant-negative manner by trapping wildtype protein in nonfunctional oligomers. Consistent with this model, the BCL11B N440K mutant interferes with BCL11A function through heterodimerization during T lymphocyte and neuronal development ^45^. However, this oligomerization-based model does not fully account for the effects of the missense variants clustered in the amino-terminal region of human BCL11A, that disrupt its dimerization ^22^. Notably, these N-terminal mutations also impair BCL11A nuclear localization, as demonstrated in HEK293T cells ^22^.

Focusing on the 16 missense mutations in BCL11B, affecting 14 different residues ^26^ (Figure S1B), we group them into four categories. The first category includes five mutations that disrupt zinc-coordinating cysteine or histidine residues in ZF5 (C826Y/R, H842P, H846D) and ZF6 (H877Y) (Figure S1C), likely destabilizing the affected ZF unit. The remaining mutants occur within the DNA-binding helix (Figure S1D-S1E) and are further divided into categories two through four. The second category includes three mutations located near the bound DNA backbone not directly contacting a DNA base, in ZF4 (C808R) and ZF5 (S836N, T840M). Substitution of small side chains (C, S, and T) with larger side chains (R, N, and M) are likely to introduce steric clashes with the DNA sugar-phosphate backbone, potentially disrupting proper ZF-DNA interactions (Figure S1F). The third category includes three mutations that alter DNA-base interacting residues in ZF3 (R472C) and ZF5 (K838T, R841L). These substitutions replace bulky, charged Arg or Lys with smaller or hydrophobic residues (C, T, and L), likely reducing DNA-binding specificity and/or affinity (Figure S1G). Consistent with this, analogous Lys-to-Thr, or Arg-to-Cys, substitutions have been characterized in BCL11A (K784T) ^34^, CTCF (K365T) ^46^, and ZBTB7A (K424T and R402C) ^47^, where the shorter Thr or Cys side chains cannot reach the DNA base and lack specific contacts, allowing the mutant to accommodate all four possible base pairs at a given position. Such broader binding specificity could also result in the BCL11B being titrated away due to increased association with the large excess of nonspecific DNA ^48^. The fourth category includes five mutations in ZF2 (N441K), ZF3 (K469N), ZF4 (N807K, N810H) and ZF5 (K838R), which alter DNA-base interacting residues by switching between polar (Asn) and positively charged residues (Lys or His), or between different positively charged residues (Lys-to-Arg) (Figure S1H), potentially shifting base recognition preferences.

Within each DNA-binding ZF unit, twelve residues separate the last Cys and the first His involved in zinc coordination. [Position numbering is relative to the first zinc-coordinating His (position 0), with residues at positions -1, -4, and -7 typically mediating DNA base recognition and underpinning the one-finger-three-base pair recognition rule ^49–51^.] The base-recognition positions are commonly occupied by larger, charged, or polar amino acids that preferentially recognize purine bases – for example, Arg/Lys/His for guanine and Asn/Gln for adenine ^52,53^. In contrast, smaller aliphatic amino acids at these positions generally permit greater variability in the recognized base pair. An invariant serine at position -5 in each ZF unit forms a cross-strand, non-specific interaction, as previously described ^54^. Based on these principles, our modeling of the third category mutations suggests that the substitution at position -1 in ZF3 (R472C) and ZF5 (R841L), as well as at position -3 in ZF5 (K838T), each of which replace bulky, charged Arg or Lys with smaller, hydrophobic residues, are likely to reduce DNA-binding specificity and/or affinity (Figure S1G). Accordingly, in subsequent analyses we focused on five mutants in the fourth category – four at position -3: N441K (ZF2), K469N (ZF3), N810H (ZF4), and K838R (ZF5), and one at position -7: N807K (ZF4), examined in the context of the isolated ZF2-3 and ZF4-6 arrays (Figure 1C). In addition, we generated a construct connecting ZF2-3 and ZF4-6 through a 22-residue linker, in place of the natural 309 residues (481-789) (Figure 1D).

### BCL11B binds the same DNA substrates as BCL11A *in vitro*

The two ZF cluster arrays (ZF2-3 and ZF4-6) in BCL11A and BCL11B function as canonical DNA-binding fingers and share high sequence identity (Figure 1B). Notably, the base-interacting residues at positions -1, -4 and -7 are identical between the two proteins, suggesting that they recognize closely related DNA sequences. Chromatin immunoprecipitation sequencing (ChIP-seq) or CUT&RUN analysis of BCL11A in HUDEP-2 cells identified a short, six-nucleotide consensus binding motif: TGnCCA (n = any nucleotide) ^55^, and the related motif TGnCC<u>w</u> (w=a or t) ^3^. Consistent with these findings, we performed *in vitro* DNA binding assays of BCL11A using four 20-bp DNA oligonucleotides (oligos) encompassing all four variants of TGnCCA ^34,35^. These four DNA duplexes were derived from BCL11A binding sites within the β-globin locus ^34^.

We next compared four proteins fragments – the isolated ZF2-3 and ZF4-6 arrays from both BCL11A and BCL11B – against the four DNA duplexes using fluorescence polarization (FP) and electrophoretic mobility shift assays (EMSA) (Figure 2). Three key observations emerged. First, corresponding ZF arrays from BCL11A and BCL11B exhibit nearly identical binding affinities: in each protein ZF4-6 binds with high affinity (K_D_ = 0.03-0.16 μM), whereas ZF2-3 shows an order of magnitude lower affinity (K_D_ = 1-3 μM). Second, among the four oligos tested, ZF4-6 from both proteins binds most strongly to the TG<u>a</u>CCA motif (K_D_ = 26-29 nM), followed by TGtCCA (K_D_ = 34-35 nM), TGgCCA (K_D_ = 58-69 nM), and TGcCCA (K_D_ = 110-160 nM). Third, two oligos each exhibited two shifted bands in EMSA assays. For the TGcCCA-containing oligo, ZF4-6 produces a first shifted band at as low a protein concentration as 5 nM (lane 4 in Figure S2), with a second, higher-molecular weight shift appearing at protein concentrations above 0.64 μM (lane 11) (Figures 2F and S2). Interestingly, the TGgCCA-containing oligo also shows a second shift with ZF2-3, despite its lower affinity (Figure 2H). Notably, both oligos (derived from the native sequences) contain partial secondary matches to the consensus motif (acCCCA or TtGCCt), which may contribute to these second binding events.

**Figure 2.**
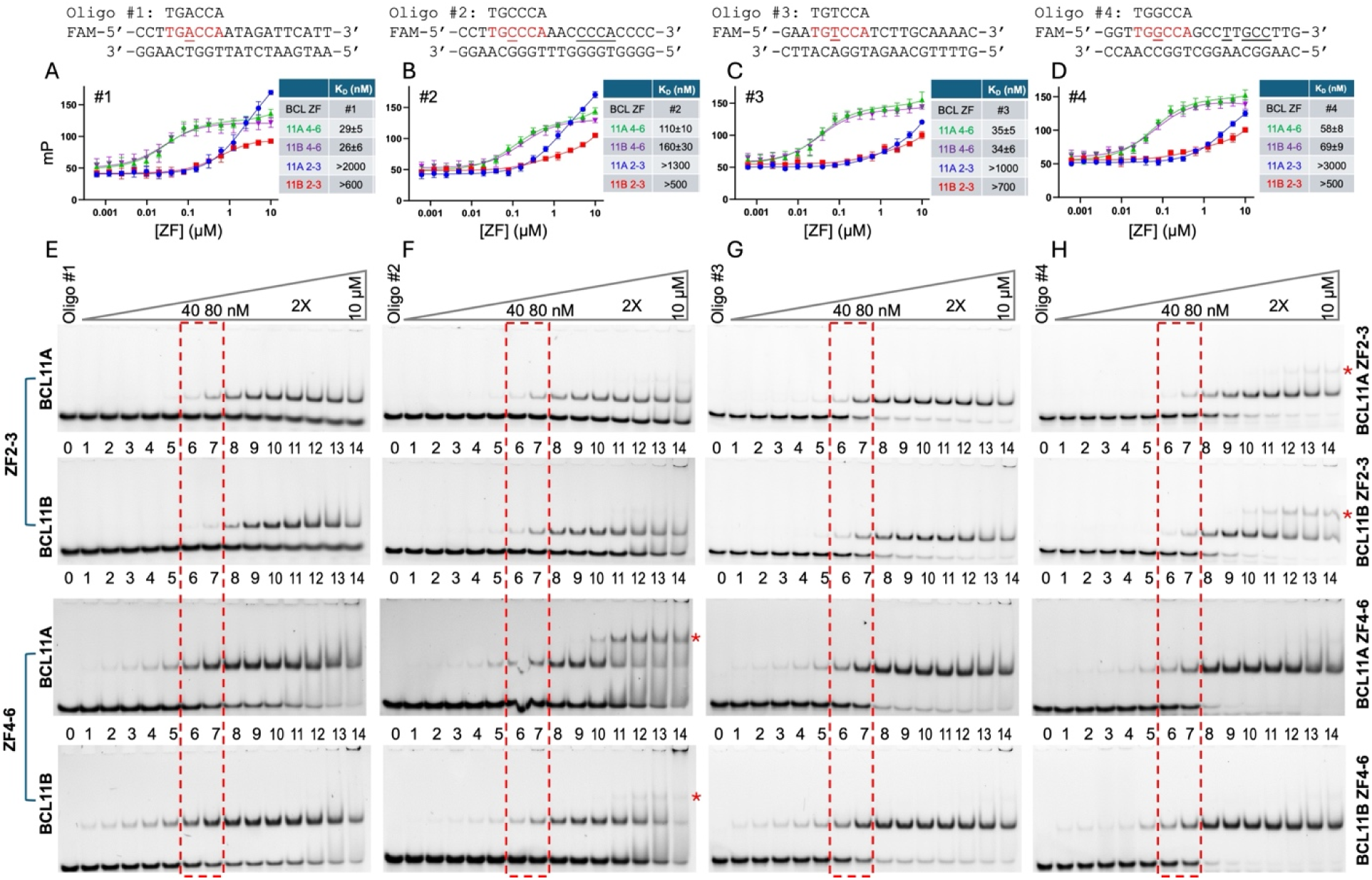
BCL11B binds the same DNA substrates as BCL11A in vitro. The same samples were used in both FP and EMSA, with the DNA duplex at 20 nM in all experiments. **(A-D)** FP assays of four oligos (sequences shown at the top of each panel) with the isolated ZF2-3 and ZF4-6 arrays from both BCL11A and BCL11B (error bars represent SEM from N = 4 independent measurements). In each case, derived K_D_ values are indicated on the right. For binding curves of ZF2-3 that did not reach saturation, the lower limit of the binding affinity was estimated. (**E-H**) EMSA of the four oligos with the corresponding ZF arrays. The red asterisks indicate the presence of a second, higher-order shifted band.

### Structures of BCL11B ZF2-3 in complex with DNA in two opposite orientations

Numerous studies have established the structural basis of DNA recognition by the BCL11A ZF4-6 using DNA substrates ranging from 12 to 20-bp and encompassing all four TGnCCA variants ^33–35,56^. In these structures, ZF4 and ZF5 make base-specific contacts within the DNA major groove, whereas ZF6 primarily engages the DNA phosphate backbone in the minor groove, contributing to overall binding affinity. We did not pursue structural analysis of the BCL11B ZF4-6 array, as the base-interacting ZF4 and ZF5 fingers are identical between BCL11A and BCL11B; sequence differences are confined to the linker between ZF5 and ZF6 and the C-terminal region following ZF6 (Figures 1B and S1A).

We recently determined structures of the BCL11<u>A</u> ZF2-3 array in complex with a TGtCCA-containing DNA in two crystal forms ^35^. Here, we extend this analysis to the BCL11<u>B</u> ZF2-3 array. Consistent with the high sequence identity between BCL11A and BCL11B at ZF2-3 – differing by only two conserved substitutions in ZF3 (Figure 1B) – screening of ∼20 oligos of varying lengths and sequences yielded crystals of BCL11B ZF2-3 bound to the same 12-bp containing a single TGtCCA motif, either with 3’-overhangs or blunt ends (Figure 3).

**Figure 3.**
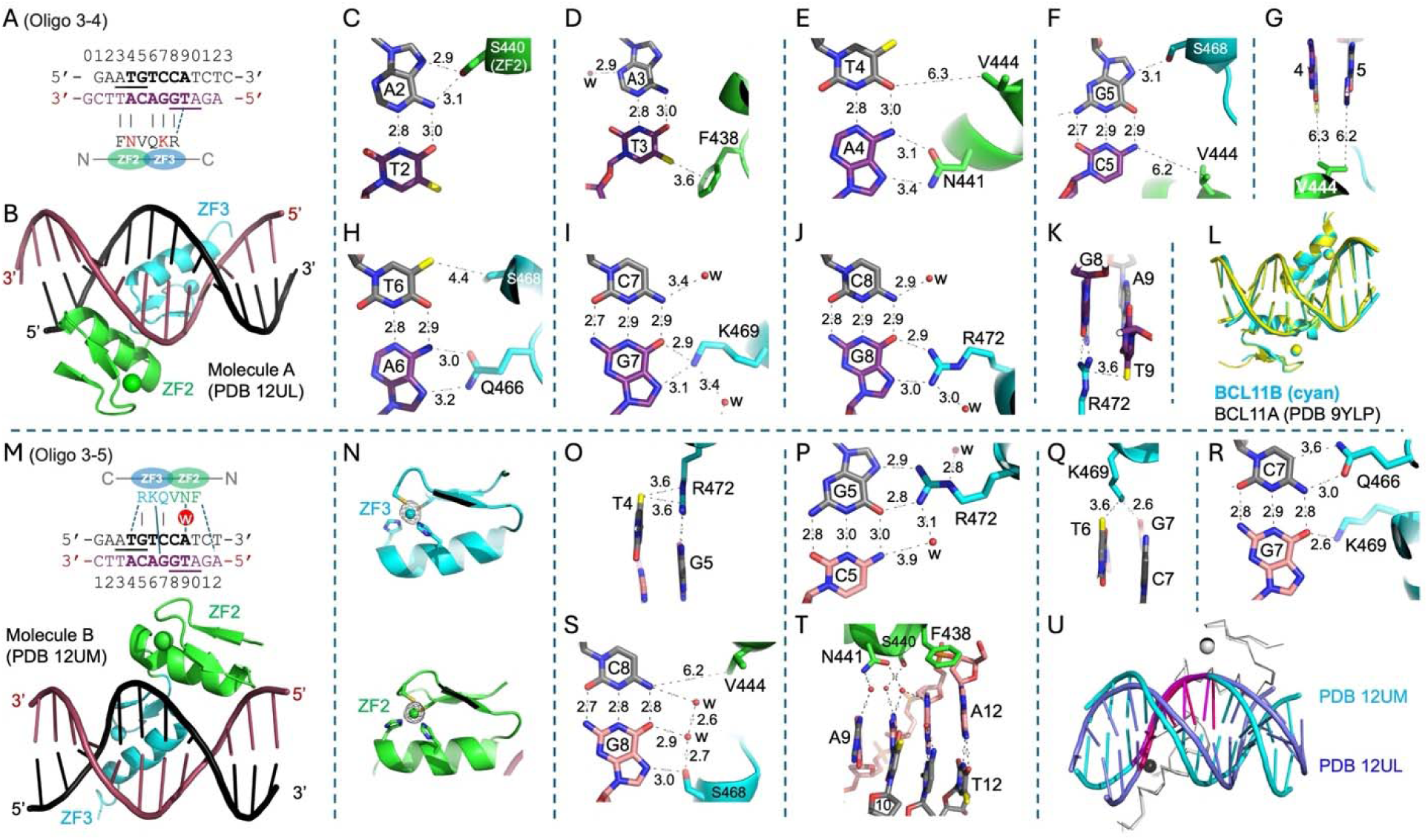
Structures of BCL11B ZF2-3 in complex with DNA in two opposite orientations. (**A-B**) Structure of ZF2-3 (molecule A) bound with a 12-bp DNA with 3’ overhangs (PDB 12UL). Base-pair numbering is shown above the DNA sequence. (**C**) S440 of ZF2 interacts with adenine A2. (**D**) F438 of ZF2 interacts with thymine T3. (**E**) N441 of ZF2 interacts with A4. (**F**) S468 of ZF3 interacts with guanine G5. (**G**) V444 of ZF2 is positioned too far from the DNA to make base contacts. (**H**) Q466 of ZF3 interacts with A6. (**I**) K469 of ZF3 interacts with G7. (**J**) R472 of ZF3 interacts with G8. (**K**) Methyl-Arg-Gua interactions among T9-R472-G8. (**L**) Superimposition of DNA-bound ZF2-3 of BCL11B (cyan) with that of BCL11A (yellow; PDB 9YLP). (**M**) Structure of ZF4-6 (molecule B) bound to a blunt-ended 12-bp DNA. (**N**) Anomalous electron densities for zinc atoms, contoured at 5.0σ above the mean, used to position each zinc finger (PDB 12UM). (**O**) Methyl-Arg-Gua interactions among T4-R472-G5 in molecule B. (**P**) R472 of molecule B interacts with G5. (**Q**) K469 of molecule B interacts with T6 and G7. (**R**) Q466 and K469 of molecule B interact with the C7:G7 base pair. (**S**) S468 of molecule B interacts with G8. (**T**) Water-mediated DNA interactions involving ZF2 of molecule B. (**U**) Superimposition of two ZF2-3 molecules (A and B) of BCL11B. The ZF3-bound four nucleotides (TGTC of the top strand and TGGA of the bottom strand), including the TpG dinucleotides, overlay well (colored magenta).

The two-finger fragment ZF2-3 (molecule A) binds within the DNA major groove and spans eight base pairs, with ZF2 contacting base pairs 2-5, while ZF3 contacting base pairs 5-9 (Figures 3A-3B). Base-pair position 5 is contacted by both fingers, with overlapping interactions from V444 of ZF2 and S468 of ZF3 (Figure 3F). We note that the direction of this ZF array (from N-to-C) runs opposite to the recognition strand (from 5’-to-3’). Base-specific interactions occur primarily on the bottom strand in Figure 3A, mediated by F438 (position -7 of ZF2) contacting the methyl group of thymine T3 (Figure 3D), N441 (−4 of ZF2) recognizing adenine A4 (Figure 3E), Q466 (−7 of ZF3) contacting A6 (Figure 3H), K469 (−4 of ZF3) recognizing guanine G7 (Figure 3I), and R472 (−1 of ZF3) contacting G8 (Figure 3J). These interactions – Asn/Gln with adenine and Lys/Arg with guanine – represent classical purine recognition via bidentate hydrogen bonds ^52,53^. In addition, S440 (ZF2) and S468 (ZF3), located at position -5 in each finger, form cross-strand, non-specific contacts with A2 and G5, respectively (Figures 3C and 3F). A hydrophobic residue, V444 (−1 of ZF2), is positioned too far from the DNA to make direct base contacts (Figures 3F and 3G). A distinctive interaction is observed for F438 of ZF2, whose aromatic ring packs against the methyl group of T3, forming a CH-π interaction (Figure 3D). Although aromatic residues are uncommon at base-interacting positions, a similar strategy is observed in ZNF410, where Tyr and Trp interact with thymine methyl groups ^57^. Finally, an additional hydrophobic interaction is observed between R472 and the methyl group of T9 (Figure 3K), consistent with the previously described methyl-Arg-Gua triad of TpG sequences ^58^. The DNA-bound ZF2-3 structure of BCL11B overlays closely with that of BCL11A (PDB 9YLP) ^35^, with a root-mean-square deviation (RMSD) of 0.425 Å across 55 pairs of C_α_ atoms (Figure 3L).

The 12-bp oligo used for crystallization exhibits pseudosymmetry, with ATG trinucleotides present on both strands, allowing R472 of ZF3 to engage a TpG dinucleotide in either orientation. Consistent with this, we observed a second crystal form in which the ZF2-3 array binds in the opposite orientation (molecule B in Figure 3M), engaging the top strand. To confirm the correct protein orientation, anomalous electron density from the zinc atoms was used to unambiguously position each zinc finger (Figure 3N). In molecule B, interactions between T4-R472-G5 on the top strand mirror those observed for T9-R472-G8 on the bottom strand in molecule A (comparing Figures 3O and 3P to 3J and 3K). K469 of ZF3 spans base pairs 6 and 7, forming van der Waals contacts with T6 (Figure 3I), while its terminal amino group forms a cross-strand hydrogen bond with G7 of the opposite strand (Figure 3R). Q466 of ZF3 also contacts C7, the base paired with G7. Similarly, S468 of ZF3 forms a cross-strand interaction with G8 (Figure 3S), analogous to the S468-G5 interaction observed in Figure 3F. Further along the DNA, ZF2 of molecule B engages in water-mediated interactions involving the polar residues S440 and N441, while F438 is positioned above the ribose ring of A12 (Figure 3T). Although the two ZF2-3 molecules (A and B) superimpose well (RMSD = 0.58 Å across 55 pairs of C_α_ atoms), the DNA conformations differ substantially, particularly at the duplex ends where protein contacts are absent (Figure 3U).

Because ZF2-3 can bind DNA in two orientations, we redesigned the 12-bp DNA sequence to be palindromic (5’-ATATGGCCATATG-3’) with a 3’ G overhang. This DNA element yielded two crystal forms, each containing three DNA duplexes stacked head-to-end within the crystallographic asymmetric unit (Figure S3). In space group *P*2_1_2_1_2_1_, three DNA-bound ZF2-3 molecules were arranged in two distinct orientations (Figure S3A). In space group *C*2, only two of three DNA duplexes were bound by ZF2-3, and these were arranged in opposite orientations (Figure S3B). In each ZF2-3 molecule, K469 and R472 of ZF3 fully engaged two consecutive guanines on one strand.

### Behavior of Missense Variants

We first examined the effects of the N441K (ZF2) and K469N (ZF3) mutations in the context of ZF2-3. FP assays were not suitable for distinguishing differences between wild-type (WT) and mutant proteins due to the relatively low binding affinity (K_D_ = 1-3 μM) and protein aggregation at the higher concentrations (> 3 μM). Therefore, binding was assessed by EMSA (Figure S4). Under the conditions tested, both single-point mutants retained the ability to bind DNA. However, they exhibited reduced binding affinity compared to WT, as the WT protein starts to induce a shift at 40-80 nM, whereas the mutants required higher concentrations (Figure S4A-B). For the oligo containing TGTCCA, the difference between the WT and the mutants was minimal (Figure S4C). For the oligo containing TGGCCA, the WT protein produced a second shifted band at higher concentrations, but this second shift was absent for the mutant proteins (Figure S4D).

Next, we characterized four mutants within ZF4-6: N807K (ZF4), N810H (ZF4), K838R (ZF5), and K838T (ZF5). Because ZF4-6 binds DNA with nanomolar affinity, this construct enabled quantitative FP comparison between WT and mutant proteins (Figure 4). Compared to WT, N807K showed a ∼2.5X reduction in affinity for oligos #1 and #2, little change for oligo #3 (1.1X), and increased affinity for oligo #4 (0.58X). The enhanced binding to oligo #4 may be explained by a favorable interaction between lysine at position 807 and a guanine (highlighted in red; Figure S5). N801H exhibited a 4-9X reduction in affinity across all four oligos. Among the substitutions, K838R had the least effect, consistent with both Lys and Arg being positively charged residues that recognize guanine. For this mutant, three oligos showed minimal changes (0.9X, 1.0X, and 1.4X relative to WT), whereas oligo #4 showed the largest affinity decrease (4.7X; Figure 4C). Finally, K838T caused the most pronounced loss of affinity, with reductions of 23X for oligo #1 and >30X for oligo#4, along with more modest decreases for oligo #2 (3.9X) and oligo #3 (2.3X). Overall, the data indicate that these mutations can result in both loss- and gain-of-function effects on DNA binding in a sequence-dependent manner.

**Figure 4.**
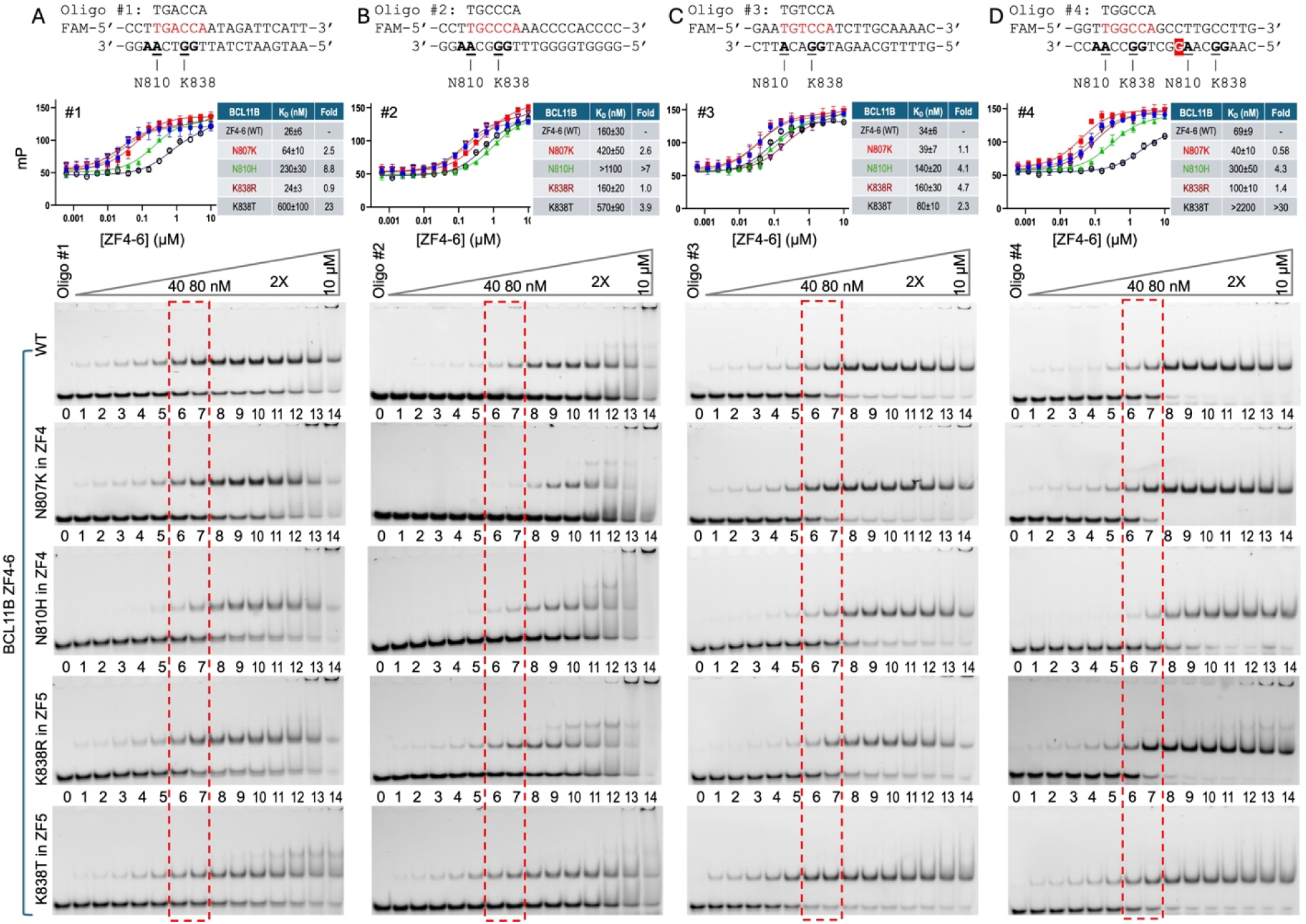
Binding affinities of mutants in ZF4-6. (A-D) Four variable DNA sequences (shown at the top of each panel). Potential base-specific interactions are indicated: **AA** contacts (N807 and N810 of ZF4) and **GG** contacts (K838 and R841 of ZF5). For each oligo (at 20 nM), FP binding curves and derived K_D_ values are shown for each protein (error bars represent SEM from N = 4 independent measurements). EMSA results for the same samples used in FP are shown for five protein variants in the following order: WT, N807K (ZF4), N810H (ZF4), K838R (ZF5), and K838T (ZF5). At the highest protein concentrations (lane 14; 10 μM), protein precipitation is frequently observed in the loading well.

### BCL11B binds regulatory elements of SHANK3

An ENCODE Project ChIP-seq dataset of eGFP-BCL11B in HEK293 cells is available (accession: ENCSR770PQN). Although this dataset does not reflect endogenous expression in a physiologically relevant cell type, Lessel *et al.* identified ∼9,000 BCL11B binding peaks ^26^. Of these, just under half (4,123; 46%) are located within ±1 kb of annotated transcription start sites. The activity of candidate *cis*-regulatory elements from three genes – *SHANK3* (<u>SH</u>3 and multiple <u>ank</u>yrin repeat domains <u>3</u>), *LTBP3* and *LTBP4* (<u>L</u>atent <u>T</u>GFβ <u>b</u>inding <u>p</u>roteins 3 and 4) (see Figure S7) – was evaluated using a luciferase assay ^26^. *SHANK3* is a large gene (∼60 kb) located on chromosome 22, encoding a 1806-residue synapse-associated protein that is implicated in a range of neurodevelopmental disorders, including intellectual disability, autism spectrum disorder, and epilepsy ^59^. Here, we focus on a BCL11B-binding peak located near exon 17 (of 23 total) and upstream of transcript ENST00000664402.3 (Figure 5A), which is derived from the remaining six exons and shows the highest median expression in the cerebellum (information from UCSC genome browser ^60^).

**Figure 5.**
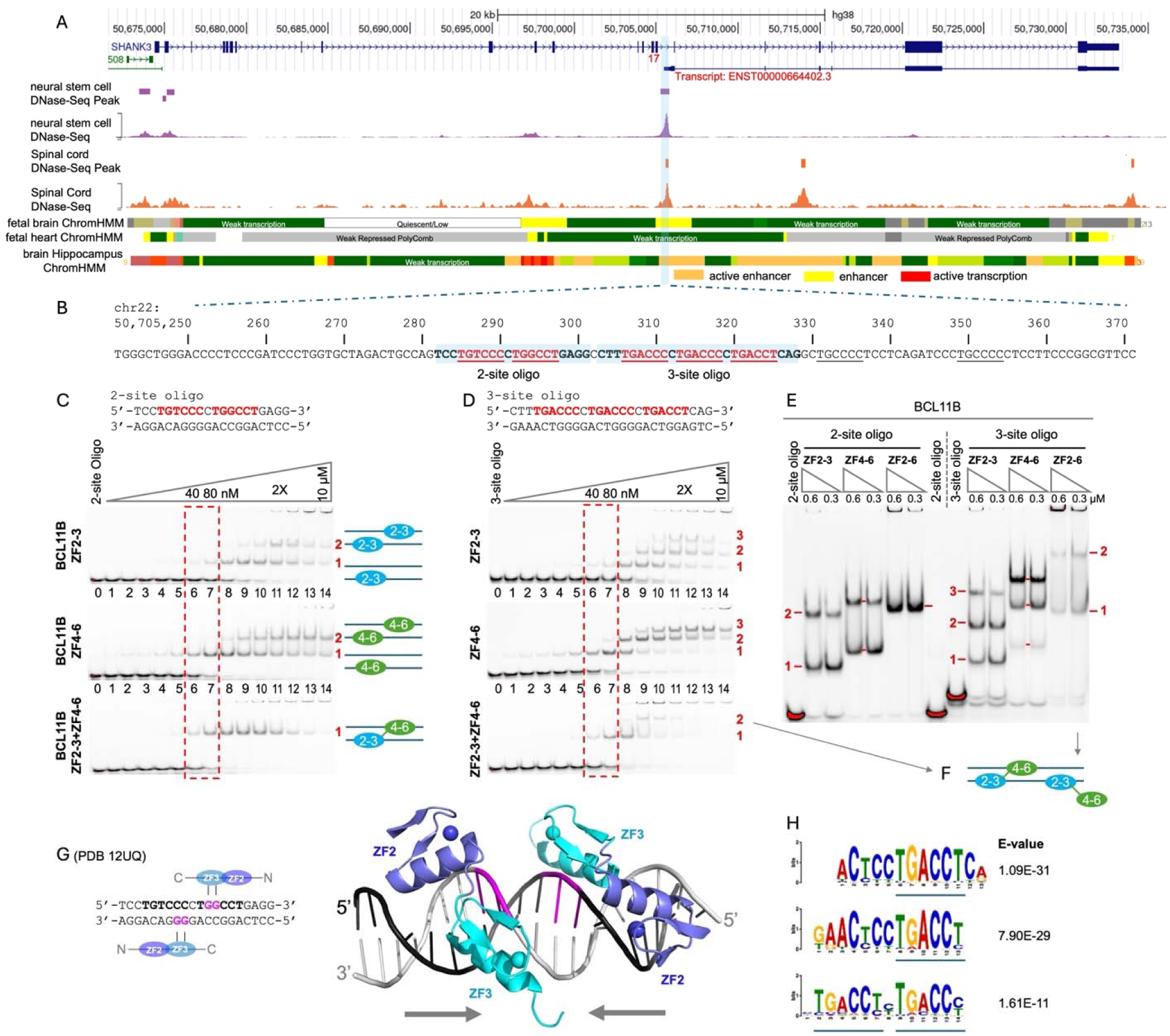
BCL11B binds the regulatory element of SHANK3. (**A**) Genome browser view of the *SHANK3* locus on chromosome 22 (from UCSC genome browser). The BCL11B binding site is located between exon 17 and the start site of transcript ENST00000664402.3. (**B**) Sequence of the regulatory element (indicated by a vertical bar in panel A), showing clusters of two or three motifs separated by a single nucleotide. This binding peak lies within an ENCODE candidate *cis*-regulatory element (EH38E3491648). (**C**) EMSA of the two-site oligo with three protein fragments, as indicated on the left. A possible binding model is shown on the right. (**D**) EMSA of three-site oligo. (**E**) EMSA of 2-site and 3-site oligos with three different protein fragments at two concentrations each. (**F**) A possible model of the three-site oligo bound by two linked arrays, in which one array (either ZF2-3 or ZF4-6) does not engage DNA specifically. (**G**) Structure of the 2-site oligo bound with two molecules of ZF2-3, arranged in opposite orientations. (**H**) Top enriched centrally oriented motifs identified by motif discovery in eGFP-BCL11B ChIP-seq peaks from HEK293 cells (see also Figure S7D).

The BCL11B binding peak covers multiple copies of the six-nucleotide motif, TG(t/g/a/c)CC(c/t), with variation at positions 3 and 6 (Figure 5B). Notably, there are clusters of two or three motifs next to each other and separated by one nucleotide. To investigate binding to this peak sequence, we synthesized a 20-bp oligo containing TGtCCc and TGgCCt motifs, and a 26-bp oligo containing three TGACC(c/t) motifs, and analyzed BCL11B binding via EMSA. For the two-site oligo, using isolated ZF arrays (ZF2-3 or ZF4-6), a single shifted band appeared at relatively low concentrations (40-80 nM) for ZF2-3 (lanes 6-7), and at even lower concentrations for ZF4-6 (lanes 4-5) (Figure 5C, top and middle panels). A second shifted band became apparent at protein concentrations above ∼0.3 μM (lanes 9-10), suggesting that the two-site oligo can accommodate two independent binding events by the isolated ZF arrays. We next tested a linked construct containing a 22-residue linker connecting ZF2-3 to ZF4-6 (Figure 1D). In this case, a single shifted band was observed, suggesting that both ZF arrays within a single polypeptide can bind simultaneously to the same oligo (Figure 5C, bottom panel).

Similarly, three binding events are observed on the three-site oligo with the isolated ZF arrays, whereas two binding events are detected with the linked ZF arrays (Figures 5D and 5E). Because only three possible binding sites are available, occupancy of a single site by either ZF2-3 or ZF4-6 within the linked construct can produce the same EMSA shift (Figure 5F). These results suggest that the high sequence similarity between ZF2-3 and ZF4-5 (Figure 1B), including key DNA-contacting residues, enables both ZF arrays to recognize similar sequences. In addition, BCL11B isoforms may form oligomers ^41–43^, potentially enhancing chromatin occupancy and transcriptional activity by allowing multiple copies of both ZF arrays to bind DNA simultaneously.

To confirm multivalent binding to the two-site oligo, we crystallized the 20-bp oligo in complex with two ZF2-3 molecules (Figure 5G). The two ZF2-3 molecules bind in opposite orientations, with one recognizing the bottom strand and the other the top strand. The key anchoring interactions involve dinucleotide guanines contacted by K469 and R472 of ZF3, consistent with those described above (Figures 3I and 3J).

Motif discovery analysis of ENCODE ChIP-seq data for eGFP-BCL11B in human HEK293 cells identified multiple closely related sequences containing a TGACC(t/c) core that exhibit strong central enrichment within peak regions (Figure 5H). These motifs likely represent variants of a shared binding preference, differing primarily in flanking sequence and motif length. Notably, one motif exhibited a tandem arrangement of the TGACC(t/c) sequence separated by a single nucleotide, as observed at the *SHANK3* binding site (Figure 5B).

Substitutions in both ZF2-3 and ZF4-6 reduced binding on both two-site and three-site oligos (Figures 6 and S6). Compared to ZF2-3 WT, the N441K and K469N mutants lose the second shifted bands in EMSA, and K469N also shows reduced binding in the first shifted band (Figure 6A). Similarly, in ZF4-6, the K838T mutant loses the second shifted band, while N807K and N810H show a delayed appearance of the second shifted band at higher protein concentrations. In contrast, K838R retains binding behavior comparable to WT ZF4-6 (Figure 6B).

**Figure 6.**
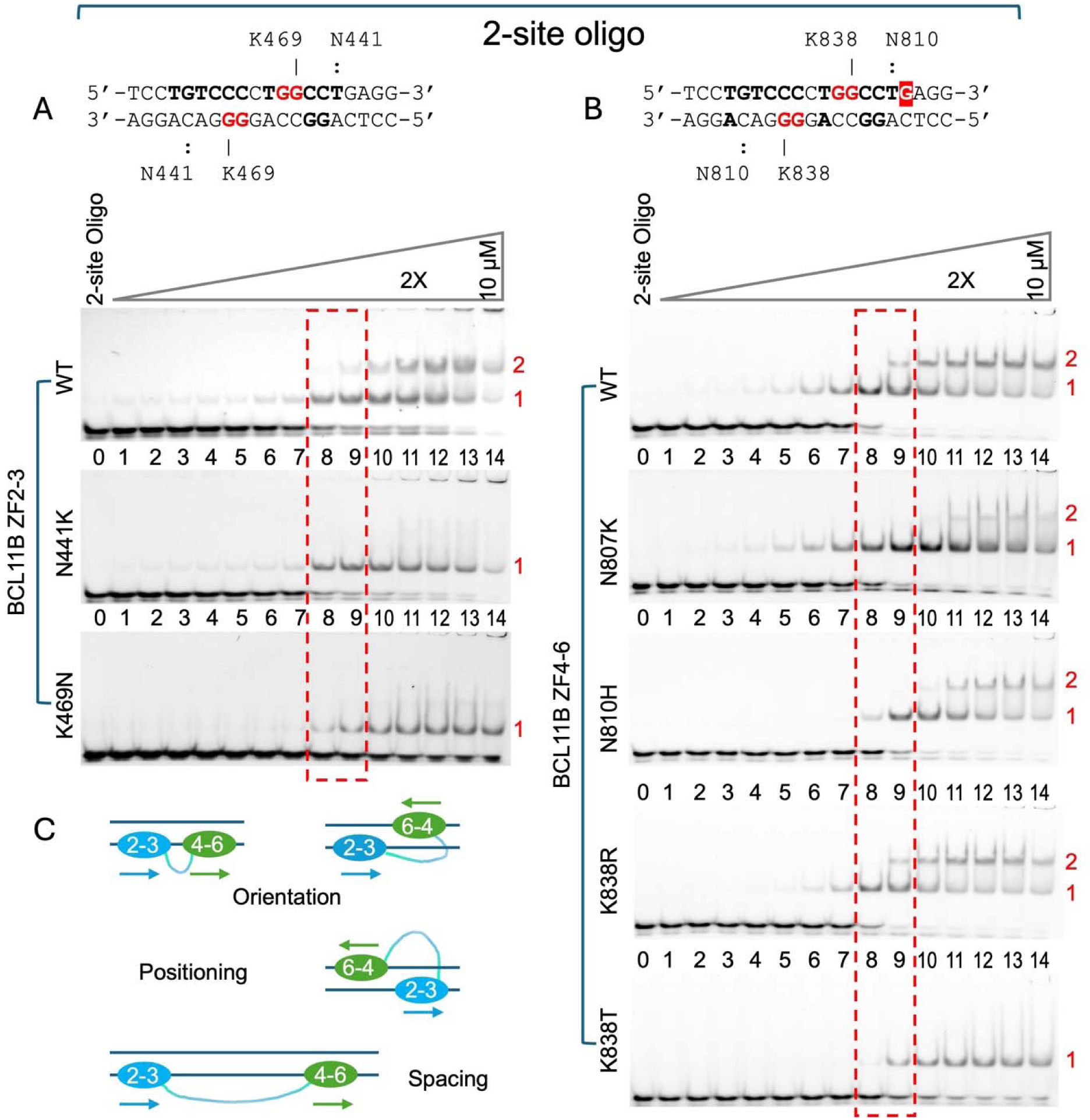
BCL11B mutants on 2-site oligo. (**A**) EMSA of ZF2-3 WT, N441K and N469K. (**B**) EMSA of ZF4-6 WT, N807K, N810H, K838R, and K838T. (**C**) Different arrangements of the ZF2-3 and ZF4-6 arrays on DNA. Arrows indicate the orientations of each ZF array from the N-to-C terminus. ZF2-3 and ZF4-6 can bind either the same stand or opposite strands, with variable spacing and positioning along the DNA.

## DISCUSSION

### Oligomerization

Dimerization is a most common mode of protein self-association ^61^, particularly among DNA-binding transcription factors (TFs), which often function as homodimers, heterodimers (with closely related family members), tetramers, or higher-order oligomers. Examples include basic leucine-zipper proteins such as AP-1 (Fos/Jun) ^62^ and C/EBP ^63,64^, basic helix-loop-helix proteins such as Myc/Max ^65^, Rel homology domain proteins such as NF-κB ^66^, nuclear receptors ^67^, and the p53 tetramer ^68^. Among these DNA-binding protein families, the majority of TF polypeptides (779) contain a single DNA binding domain (DBD), whereas only a minority (79, or 10%) contains more than two homotypic DBDs (Lambert, 2018 #4211}. Thus, oligomerization provides an effective mechanism to increase the number of DBDs available for target recognition. In BCL11B, oligomerization – through homo- or hetero-association with BCL11A – could increase the number of DBDs from two to four or even eight.

### C2H2 ZF proteins

C2H2 zinc finger proteins, numbering ∼700, constitute approximately half of ∼1500 annotated sequence-specific DNA-binding proteins in the human genome ^69,70^. However, the number of fingers per protein varies widely, ranging from ∼3 to 35 fingers, with a mode of around 11–13 fingers ^58^. Based on “one-finger-three-base” rule of recognition ^49–51^, a three-finger array can recognize ∼9 base pairs, whereas an 11-finger array can, in principle, recognize ∼33 base pairs. Accordingly, an 11-finger array of CTCF can bind thousands of diverse genomic sites (ranging from 12-15 bp to 40 bp) ^46,71–76^, whereas another 11-finger protein, ZFP568, recognizes a single specific site of 24-bp ^77,78^. Historically, structural and functional studies of ZF-DNA recognition have focused on three-finger proteins such as early growth response 1 (EGR1, also known as Zif268) ^79–85^, which also enabled the development of programmable ZF-nucleases for sequence-specific genome engineering ^86^. As shown in BCL11B, DNA-binding specificity depends on the amino acids at the base-contacting positions -1, -4 and -7 within each finger (where 0 is the position of the first Zn-coordinating His), and two-finger array of ZF2-3 recognizes a six-bp duplex.

### A combination of high- and low-affinity DNA binding

BCL11A and BCL11B contain two short ZF arrays, ZF2-3 and ZF4-6, separated by ∼300 residues (Figure 1A). These two arrays each function as homotypic DBDs that recognizing similar sequences, as seven of the eight residues involved in direct DNA base contacts are identical between ZF2-3 and ZF4-5 (Figure 1B). Although ZF6 does not directly contact DNA bases ^33,34^, its presence strengthens DNA binding, making the three-finger array (ZF4-6) up to ∼100-fold stronger than the two-finger array (ZF2-3). Importantly, several studies have shown that low-affinity DBDs provide a distinct functional contribution that cannot be replaced by high-affinity DBDs ^87–91^. A notable example is the master myeloid transcription factor PU.1, which binds DNA across a wide range of affinities, spanning approximately three orders of magnitude in dissociation constants ^92^. In the case of BCL11B, a germ line missense mutation (N441K) occurs in ZF2 of the low-affinity array ZF2-3 ^45^. The mutant acts in a dominant-negative manner, likely through the formation of a nonfunctional or dysregulated dimers ^43^, resulting in reduced binding to canonical target sites as well as aberrant binding to novel DNA sequences not recognized by the wild-type protein ^45^.

In regions bound by multiple DBDs, short DNA sequence motifs often occur in clusters. The number and organization of these clustered motifs, together with the number of DBDs, contribute to the complexity of gene regulation and may have coevolved to maintain structural integrity and functional specificity ^93,94^. One example examined here is the *SHANK3* regulatory element: within an ∼80-nt stretch in the center of BCl11B binding peak, there are seven copies of a six-nucleotide motif, TG(t/g/a/c)CC(c/t) (Figure 5B), five of which are separated by only a single nucleotide. This dense clustering coincides with a DNase-seq peak in neural stem cells and spinal cord, consistent with an accessible enhancer region (Figure 5A), and is characteristic of regulatory regions with high motifs density ^95^. The downstream transcript ENST00000664402.3 encodes a 1112-residue protein (UnitProt A0A590UJL3) that is predicted by AlphaFold to be largely disordered, except for a C-terminal ∼70-residue sterile alpha motif (SAM) domain. This domain self-associates and forms polymers that contribute to postsynaptic density assembly at the postsynaptic membrane ^96^.

### Orientation, spacing and positioning of a bipartite DNA binding domain

The two ZF arrays – low-affinity ZF2-3 and high affinity ZF4-6 – can be considered as a fused heterodimer in which the two “subunits” are covalently linked by a long ∼300-residue linker (Figure 1A). This extended linker likely provides flexibility in how the two arrays engage DNA, suggesting that there is no fixed arrangement of ZF2-3 and ZF4-6 on the DNA (Figure 6C). Structural studies of ZF2-3 of BCL11B (this work), together with ZF4-6 of BCL11A ^35^, show that both arrays can recognize diverse TG(n)CC(c/t/a) sequences and bind DNA in either orientation and on either strand. The most conserved interactions involve guanine dinucleotides contacted by Arg and Lys residues at positions -1 and -4 of ZF3 and ZF5. Together, these observations indicate that the orientations, spacing, and positioning of the ZF2-3 and ZF4-6 arrays – connected as they are by a long, flexible linker – can vary substantially with respect to one another on DNA (Figure 6C).

### Three potential layers of regulation

Additional mechanism(s), beyond the intrinsic sequence specificity of ZF arrays, likely contribute to the regulation of BCL11B DNA binding. Here, we outline three potential layers of regulation that were not directly addressed in this study. The long, flexible linker connecting ZF2-4 and ZF4-6, is enriched in serine/threonine residues, acidic residues (aspartate/glutamate), and structural residues (glycine/proline) (Figures 1A and S1A). Serine and threonine residues are potential sites of phosphorylation, a common regulatory mechanism that introduces negatively charged phosphate groups and links signaling pathways to gene expression control ^97^. Phosphorylation can modulate multiple aspects of TF function, including subcellular localization, protein stability, protein-protein interactions, and DNA binding ^98,99^. In particular, phosphorylation near a DNA-binding domain introduces negative charge, which is generally incompatible with efficient binding to the polyanionic DNA backbone ^100–103^. In primary mouse thymocytes, upon T-cell receptor (TCR) stimulation, MAP kinase- and PKC-mediated phosphorylation in Bcl11b, as well as SUMOylation, have been identified by mass spectrometry ^104–106^. Across multiple studies, 25 ^104^, 18 ^107^, and 28 ^108^ serine/threonine residues (spanning S95 to S779) have been reported to be phosphorylated in Bcl11b. Notably, a mouse model carrying 23 S/T-to-Ala substitutions in Bcl11b shows normal primary T cell development in the thymus and maintenance of peripheral T cells ^108^, suggesting that global phosphorylation of Bcl11b is dispensable for T cell differentiation. However, reduced levels of the mutant protein, together with the retention of five phosphorylation sites – including two mutant-specific sites with 100% probability of being phosphorylated (S487 near ZF2-3 and S725 near ZF4-6) ^108^ and conserved in human BCL11B (Figure S1A) – indicate that specific phosphorylation events may fine-tune Bcl11b activity in a context-dependent manner.

A stretch of negatively charged Asp and Glu residues – 13 in BCL11B and 21 in BCL11A (boxed as red E in Figure 1A) – may structurally and electrostatically mimic DNA, as observed in known DNA-mimicry proteins ^109–111^. These DNA-mimicking proteins use negatively charged carboxyl side chains to emulate DNA phosphate groups and modulate the activity of diverse DNA-binding proteins, including BRCA2 ^112^, CRISPR-Cas9 ^113^, NF-κB ^114^, HIC2 ^115^, and ZNF410 ^116^. Both HIC2 and ZNF410 are ZF transcription factors containing five-finger DBDs. In ZNF410, an acidic and S/T-rich loop precedes the DBD, whereas in HIC2, an insertion enriched in negatively charged and S/T/Y residues lies between the first two fingers. In both cases, the full-length protein ^116^ or larger fragments containing the acidic region ^115^ reduce DNA binding, consistent with a DNA-mimicry mechanism.

Glycine and proline residues play important roles in protein folding, with glycine providing flexibility and proline imposing conformational constraints. In BCL11B, these amino acids are enriched within residues 568-680, where they comprise ∼37% of the sequence (Figure 1A and S1A). Such Gly- and Pro-rich regions are often intrinsically disordered (as predicted by AlphaFold ^117^) and can promote liquid-liquid phase separation (LLPS), a process that contributes to the formation of transcriptional condensates ^118–122^; though LLPS-independent mechanisms of intrinsically disordered regions modulating DNA specificity on chromatin have also been proposed ^123^. In the context of BCL11B, this region may facilitate the assembly of high-order regulatory complexes at chromatin, including the <u>nu</u>cleosome <u>r</u>emodeling and histone <u>d</u>eacetylase (NuRD) complex ^105,124–127^, thereby enhancing or modulating DNA binding and transcriptional regulation.

In summary, a surprising amount remains to be learned about the activities of the BCL11A and BCL11B, considering the fundamental roles they play in development of blood, brain, and adaptive immunity. Here, we contribute to understanding their DNA interactions, but other aspects of their regulatory mechanisms deserve further study.

## Methods

### BCL11A and BCL11B plasmid constructs

Constructs used in this study are summarized in Figure 1E. BCL11A ZF2-3 (pXC2394) and ZF4-6 (pXC2391) plasmids were described previously ^34^ and were used for comparative analysis. The human full-length BCL11B construct pKLV-EF1a-mCherryBCL11B-W was a gift from Kosuke Yusa (Addgene plasmid #208758) ^128^. BCL11B fragments encoding ZF2-3 (pXC2442), ZF4-6 (pXC2437), and ZF2-6 (pXC2461) were generated by overlapping PCR and cloned into the pGEX-6P-1 vector. These constructs contain an N-terminal glutathione S-transferase (GST) tag followed by a human rhinovirus (HRV) type–14 3C protease cleavage site (LEVLFQGP). Point mutants of BCL11B in ZF2-3 – N441K (pXC2443) and K469N (pXC2444) – and in ZF4-6 – N807K (pXC2438), N810H (pXC2439), K838R (pXC2440), and K838T (pXC2441) – were generated by site-directed mutagenesis and confirmed by sequencing.

### BCL11B ZF arrays expression and purification

All wild-type and mutant plasmids encoding BCL11B ZF2-3 and ZF4-6 were transformed into *Escherichia coli* strain BL21(DE3) Gold+ cells for protein expression. Cultures were grown in LB medium supplemented with 50 µg/mL ampicillin at 28 °C to an OD_600_ of 0.7-0.8, ZnCl_2_ was then added to a final concentration of 25 µM, and the temperature was reduced to 15 °C. After 30 min, protein expression was induced by 1 mM isopropyl β-D-1-thiogalactopyranoside (IPTG), and cultures were incubated overnight (∼16 h).

Cells were harvested and resuspended in the HEPES buffer [20 mM HEPES. pH 7.5, 500 mM NaCl, 5% glycerol, and 0.5 mM tris(2–carboxyethyl)phosphine (TCEP)] containing 500 mM NaCl. Cells were lysed by sonication (60% amplitude; 3s on and 6s off) for two cycles of 3 min and 2 min, respectively, using a QSONICA sonicator. Lysates were clarified by centrifugation at 74,766 x g for 1 h at 4 °C.

Cleared lysates from a 500 mL culture were incubated with 2 mL GST resin for 2 h at 4 °C with gentle stirring. The resin was washed three times with 50 mL of the HEPES buffer containing 300 mM NaCl. GST tags were removed by on–resin cleavage using 80 µL PreScission protease (190 µM stock; purified in house) in 60 mL HEPES buffer at 4 °C with overnight incubation under stirring. Eluted proteins were collected and the resin was further washed three times with 20 mL wash buffer.

Proteins were purified by cation-exchange chromatography on a 5-mL HiTrap SP HP column (Cytiva) equilibrated with the TRIS buffer (20 mM Tris–HCl, pH 7.5, 300 mM NaCl, 5% glycerol, and 0.5 mM TCEP) containing 300 mM NaCl. Proteins were eluted using a linear gradient of 300–1000 mM NaCl over 10 column volumes. Peak fractions were pooled, concentrated, flash-frozen in liquid nitrogen, and stored at -80 °C for later use.

For selected BCL11B mutants, pXC2441 (ZF4-6 K838T) and pXC2444 (ZF2-3 K469N), showing substantial DNA contamination, samples were pooled from the SP column fractions and were further purified by size-exclusion chromatography on a HiLoad 16/60 Superdex S75 column equilibrated with the TRIS buffer containing 500 mM NaCl. Final peak fractions were pooled, concentrated, flash-frozen, and stored at −80 °C until use.

### Fluorescence polarization-based DNA binding assay

A 40 µL reaction mixture was prepared containing 20 nM FAM-labeled DNA duplex and varying concentrations of BCL11A or BCL11B protein in the TRIS buffer with 150 mM NaCl. Samples were incubated on ice for 20 min prior to fluorescence polarization (FP) measurement ^129^ using a Synergy Neo2 multimode reader (BioTek). FP values were fit to a one-site binding model with a nonspecific component: [mP] = [maximum mP] × [C] / (K_D_ + [C]) + [baseline mP] + NS × [C], where [mP] is the measured milli-polarization signal, [C] is the protein concentration, and NS represents the slope of the linear nonspecific binding contribution. Data were analyzed using GraphPad Prism 10.

### Electrophoretic mobility shift assay (EMSA)

Following the fluorescence polarization assay, 10 µL of each reaction mixture was analyzed on an 8% native polyacrylamide gel run in ice-cold 0.5× TBE buffer at 150 V for 25 min. Gels were imaged using Bio-RAD ChemidocTM MP Imaging system.

### Crystallography

For crystallization, purified BCL11B ZF2–3 (WT) at ∼1 mM was mixed with 12-bp DNA at a 1:1.1 molar ratio in the TRIS buffer containing 100 mM NaCl. Oligonucleotides (Table S1) were purchased from Integrated DNA Technologies (IDT) and annealed in 50 mM NaCl and 20 mM Tris-HCl pH 7.5.

For the 2-site DNA oligonucleotide (Figure 5), purified BCL11B ZF2-3 at 1 mM was mixed with DNA to a final concentration of 0.55 mM in the TRIS buffer containing 150 mM NaCl. The protein-DNA complex was incubated at room temperature for >30 min prior to crystallization.

Crystallization screening was performed using sitting-drop vapor diffusion with an Art Robbins Gryphon robot. Drops (0.2 µL protein-DNA complex + 0.2 µL reservoir) were incubated at ∼19 °C. Crystals typically appeared within 3-7 days and were flash-frozen in liquid nitrogen after brief soaking in reservoir solution supplemented with 20% ethylene glycol when needed. Crystallization conditions are shown in Table S1.

Diffraction data were collected at 100 K at beamline 17-ID-1 (AMX) at the National Synchrotron Light Source II (NSLS-II), Brookhaven National Laboratory ^130^. Data were processed using autoPROC toolbox ^131^, which employs XDS ^132^ for data reduction. Intensities from images representing 360° of crystal rotation were merged using POINTLESS and AIMLESS ^133^.

For three datasets with severe anisotropy (PDB IDs 12UL, 12UO and 12UP), as indicated by PHENIX Xtriage ^134^, STARANISO software ^135^ was used for anisotropic truncation and amplitude estimation. In such cases, resolution cutoffs were adjusted to ensure reasonable completeness and consistent R-factors across resolution shells. PHEXIX was then used for molecular replacement, map generation, and refinement ^136^. X-ray data collection and refinement statistics were summarized in Table S2. The PyMOL molecular Graphics System (Version 3.1.4, Schrödinger, LLC) was used for structural display.

### BCL11B ChIP-seq motif discovery and analysis

The ChIP-seq dataset ENCFF860DSJ was obtained from NCBI GEO (GSE92041). This dataset corresponds to the ENCODE experiment ENCSR770PQN (eGFP-BCL11B ChIP-seq in human HEK293 cells) and represents an optimal IDR-filtered peak set derived from two biological replicates processed using the ENCODE ChIP-seq pipeline ^36^. Peak regions were deduplicated, yielding 12,550 unique peaks. For motif discovery analysis, 300 bp sequences centered on peak summits were extracted. Motif discovery was performed using XSTREME (MEME Suite 5.5.5) ^137^ with a first-order Markov background model, allowing any number of motif occurrences per sequence. Motifs were ranked based on enrichment significance and central enrichment within the input sequences, and selected motifs were used for downstream analysis.

Counts of TGNCCN motifs were generated as previously described ^34^. Briefly, peak sequences (in hg38 genome assembly) were extracted using the Fasta and SeqIO modules from Pyfaidx and Biopython, respectively. Occurrences of TGNCCN motifs and their reverse complements were programmatically counted to obtain observed motif frequencies.

A first-order Markov background model was constructed from the full set of peak sequences using MEME Suite. Probabilities for all possible 6-mer sequences were derived from this model. For each peak, expected motif counts were calculated by multiplying the number of valid 6-mer windows (peak length − 5) by the corresponding motif probability. Forward and reverse-complement probabilities were combined for each motif, with palindromic motifs counted only once. Enrichment was quantified as log₂(observed/expected), with a pseudocount (ε = 1×10⁻⁶) applied to stabilize ratios. Whole-dataset enrichment values were obtained by summing observed and expected counts across all peaks.

## Statistics and reproducibility

X-ray crystallographic data were quantitatively collected and processed. Structure refinements were performed using PHENIX Refine, with 5% reflections randomly selected for cross-validation by R-free values. Data collection and refinement statistics are summarized in Supplementary Table S2. Structure quality was monitored throughout refinements and validated using the PDB validation server. Details of statistics analyses for binding experiments are provided in the legends for Figures 2 and 4 and Supplementary Figures S2 and S5.

## Data availability

The coordinates and structure factor files of the X-ray structures of the BCL11B ZF2-3 in complexes with DNA have been deposited to PDB and released under accession numbers 12UL for oligo-34 duplex in space group *P*4_1_2_1_2, 12UM for oligo-35 in space group *P*6_5_, 12UN for oligo-43 in space group *P*2_1_2_1_2_1_, 12UO for oligo-43 in space group *C*2, 12UP for oligo-42 in space group *P*1, and 12UQ for the 20-bp 2-site oligo in space group *C*2.

## Acknowledgements

The work is supported by the grants R35GM134744 (to X.C.) and R01DK132286 (to Y.H. and X.C.) from the U.S. National Institutes of Health. M.Y. is supported by a training fellowship from the Gulf Coast Consortia on the National Library of Medicine Training Program in Biomedical Informatics and Data Science (T15LM007093). M.D.M. is an undergraduate at the Rice University and F.A.K. is an undergraduate at the Texas A&M university, both were supported by the CATALYST Summer Research Training Program. M.D.M. was supported by CPRIT Training Program (RP210028) and UPWARDS (Undergraduate Students Working Towards Research in Science) Training Program (R25CA240137). F.A.K. was supported by the MD Anderson Summer Program in Cancer Research (R25CA181004). We thank Dr. Xiangpeng Kong (New York University) for assistance of access to 17-ID-1 beamtime and the beamline scientists of the National Synchrotron Light Source (NSLS) II, Brookhaven National Laboratory. The use of NSLS II resources 17-ID-1 was supported by the U.S. Department of Energy under contract DE-SC0012704.

## Author Contributions

J.L. and J.Z. performed expression, purification, DNA binding assays, mutagenesis and crystallization; J.R.H. performed X-ray diffraction experiments. M.Y. analyzed ENCODE ChIP-seq data. M.D.M. and F.A.K., under the guidance of J.L. and J.Z., performed FP and EMSA binding assays. Xiaotian Z. and Y.H. provided advice on BCL11B biology and participated in discussion. R.M.B. participated in discussions and in preparing the manuscript. Xing Z. and X.C. designed and organized the scale of the study. X.C. is a CPRIT scholar in cancer research.

