## Supplementary for "Bipartite DNA binding domain of transcription factor BCL11B binds clustered short DNA sequence motifs"

Figures: S1-S7

Tables: S1-S2

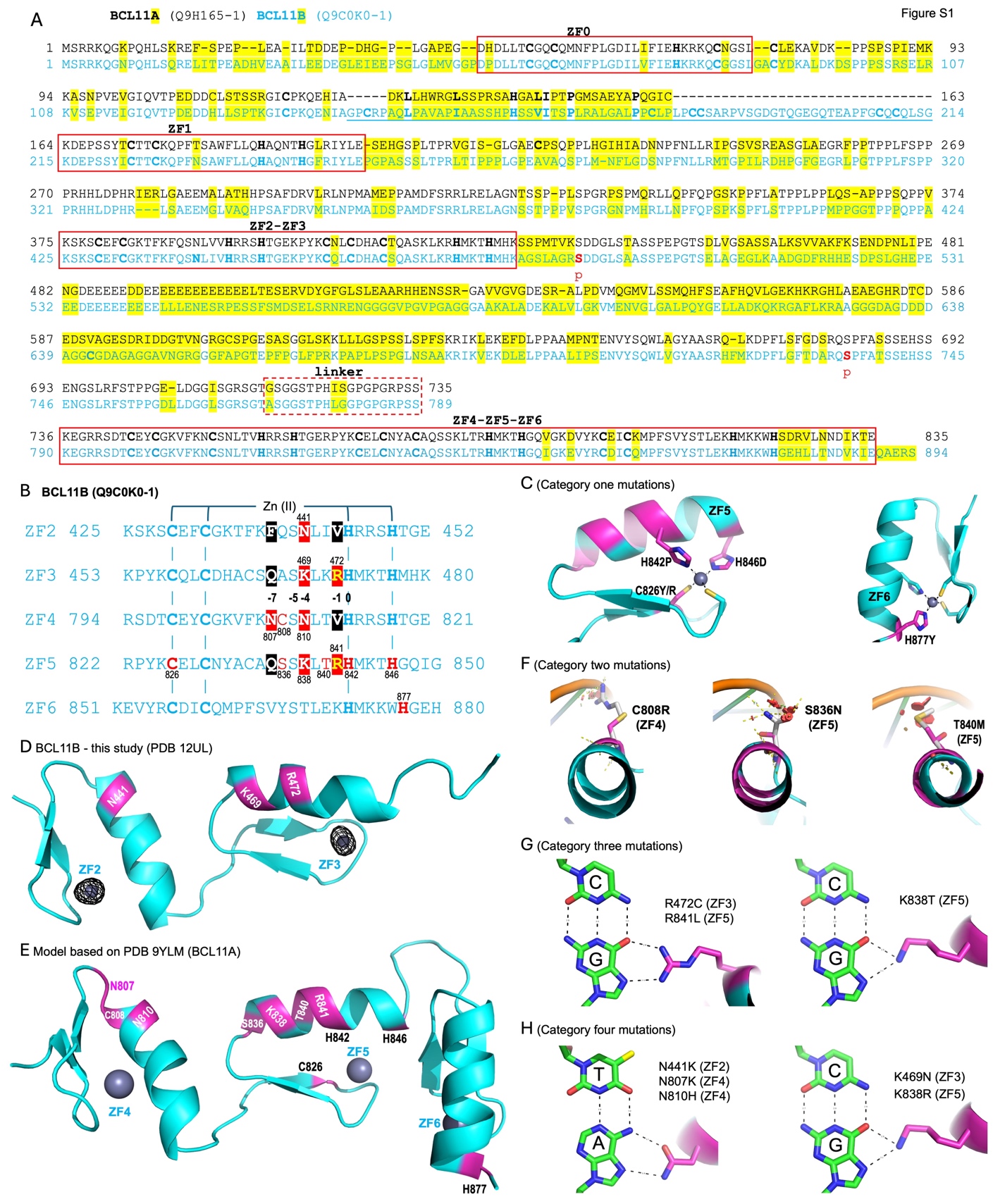

**Figure S1,** related to Figure 1, (**A**) **Sequence alignment of human BCL11A** (UniProt Q9H165-1) **and BCL11B** (UniProt Q9C0K0-1). Comparing the two sequences, BCL11A contains seventeen small deletions of 1-2 residues and one insertion of 3 residues, while BCL11B includes a 36-residue insertion (underlined) between ZF0 and ZF1 and an additional 5 residues at the C-terminus. Sequence variations between the two proteins are highlighted in yellow. Related to Figure 1B, conserved ZF regions (ZF0, ZF1, ZF2-ZF3, and ZF4-ZF5-ZF6) are labeled above the sequences and boxed in red, while the linker is indicated with a dashed box. Potential phosphorylation sites, labeled as p under S488 and S735, are derived from S487 and S725 of mouse Bcl11b (Q99PV8-1) ^1^. (**B**-**H**) **Models of 16 missense mutations in BCL11B affecting 14 residues in ZF2 to ZF6** (highlighted in red). (**B**) Sequence alignment of BCL11B ZF2-3 and ZF4-6. The zinc-coordinating C2H2 are in bold, and the DNA-base interacting residues at positions -1, -4, and -7 are boxed. Missense mutations are shown in red letters or in boxed red background. The corresponding residue numbers are shown either above or below the sequence. ZF5 contains the highest density, with nine mutations affecting seven residues. (**C**) Category one mutations that disrupt zinc-coordinating cysteine or histidine residues in ZF5 and ZF6. (**D-E**) Structural mapping of mutations onto the ZF2-3 and ZF4-6 arrays. The zinc anomalous single in panel D is contoured at 5σ above the mean. (**F**) Category two mutations near the DNA backbone involving substitutions from small to larger side chains, likely introducing steric clashes with the DNA phosphate backbone. (**G**) Category three mutations at DNA-base interacting positions that replace bulky, charged Arg or Lys with smaller or hydrophobic residues, likely reducing DNA-binding specificity and/or affinity. (**H**) Category four mutations at DNA-base interacting positions that switch between polar (Asn) and positively charged residues (Lys or His), or between different positively charged residues (Lys-to-Arg), potentially altering base recognition preferences.

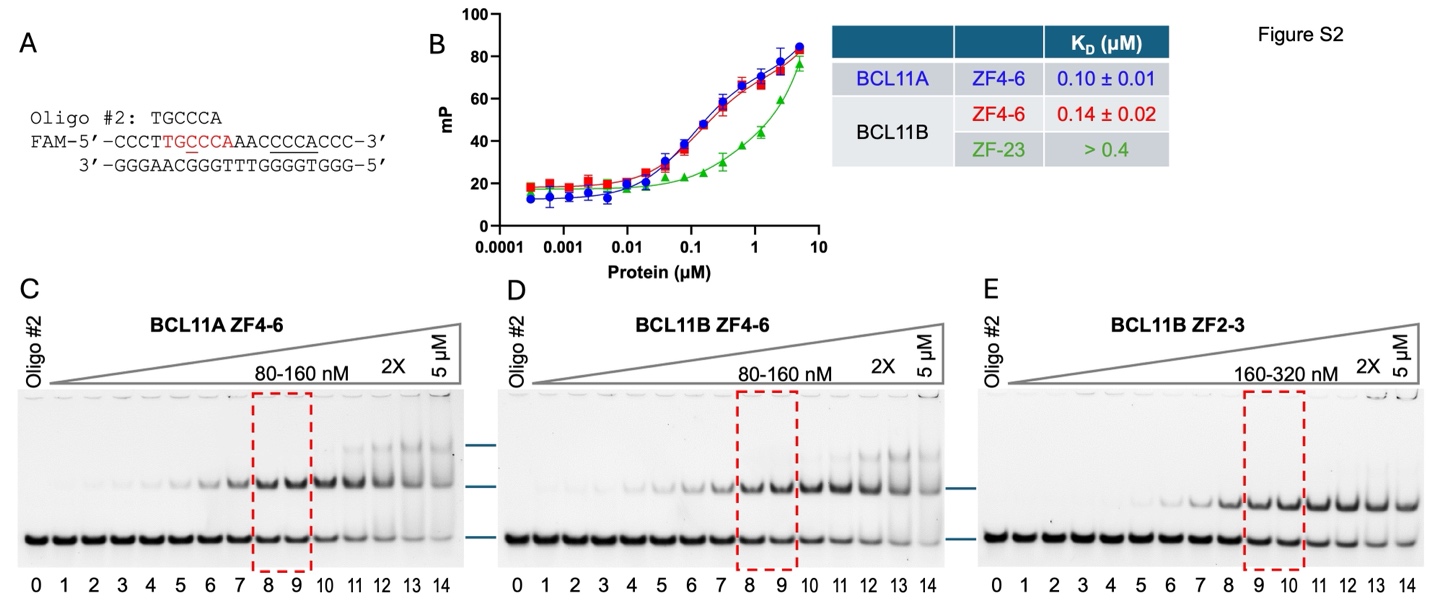

**Figure S2**, related to Figure 2, Comparison of BCL11A and BCL11B binding to TGcCCT- oligonucleotide. (**A**) Sequence of the 20-bp TGcCCT-containing oligo with a partial second match (CCCA) to the binding motif. (**B**) FP assays of BCL11A (ZF4-6) and BCL11B (ZF4-6 and ZF2-3) (Error bars represent SEM from N = 2 independent measurements). Derived K_D_ values are indicated on the right. In agreement with previous observations for BCL11A ZF2-3 ^2,3^, the two-finger ZF2-3 array of BCL11B exhibits 3-4-fold weaker binding (K_D_=0.4 μM). (**C-D**) EMSA of ZF4-6 binding. Lane 0 is DNA alone, and the other lanes have 2X series dilution starting from 5 μM at lane 14. The first shifted band appears as early as at 5 nM (lane 4), and a second, higher-order shift becomes evident at protein concentrations above 0.64 μM (lane 11). (**E**) EMSA of BCL11B ZF2-3. Consistent with its lower binding affinity, the first shift occurs at 20 nM (lane 6), and no super-shifted bands are observed at higher protein concentrations.

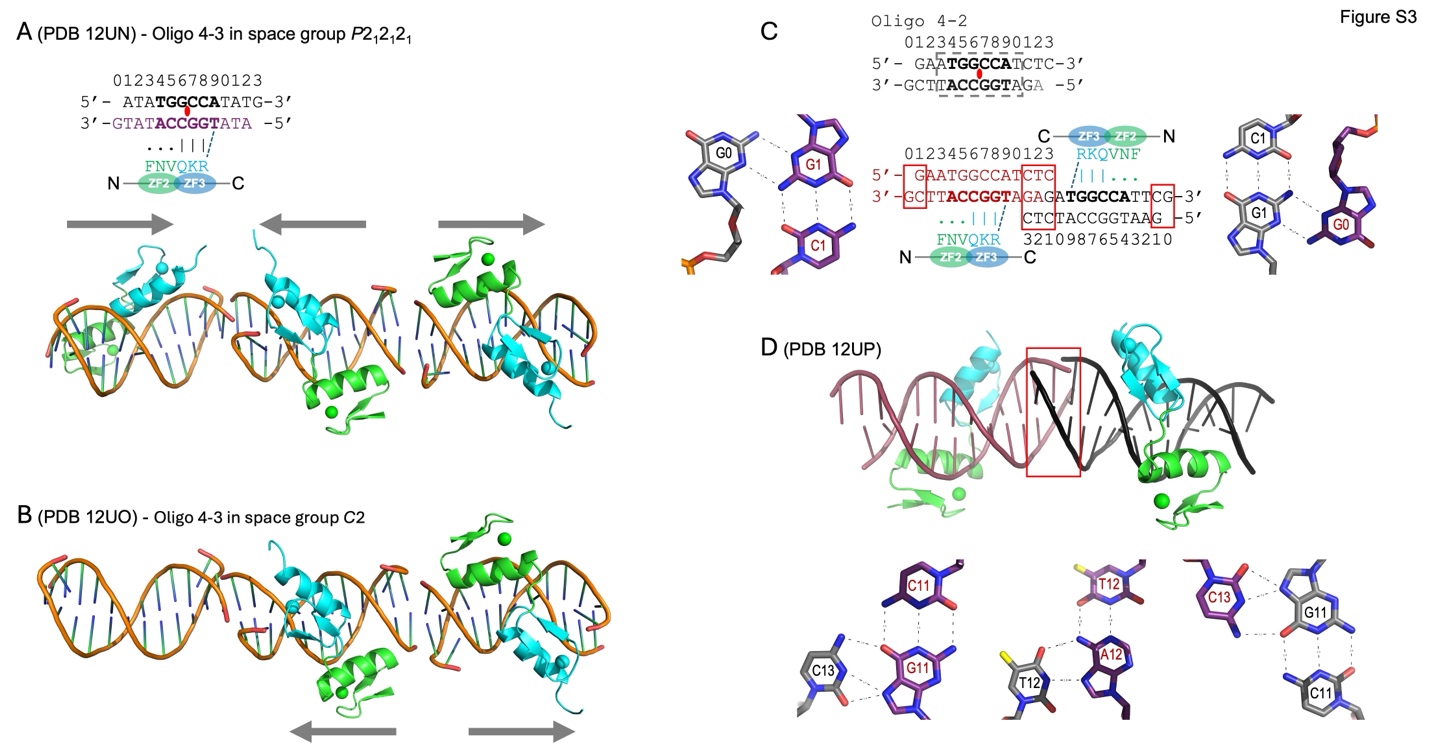

**Figure S3**, related to Figure 3, **BCL11B ZF2-3 bound to a palindromic DNA sequence**. (**A**) In space group *P*2_1_2_1_2_1_ (PDB 12UN), three protein-DNA complexes are present; arrows indicate the N-to-C orientation of ZF2-3. (**B**) In space group *C*2 (PDB 12UO), only two of three DNA duplexes are bound by ZF2-3 and are arranged in opposite orientation. (**C**). In a third structure, crystallized in space group *P*1 (PDB 12UP), two DNA duplexes pack head-to-head. The central 8-bp sequence is symmetric whereas the terminal base pairs are not. (**D**) The junction between the duplexes adopts a three-stranded conformation.

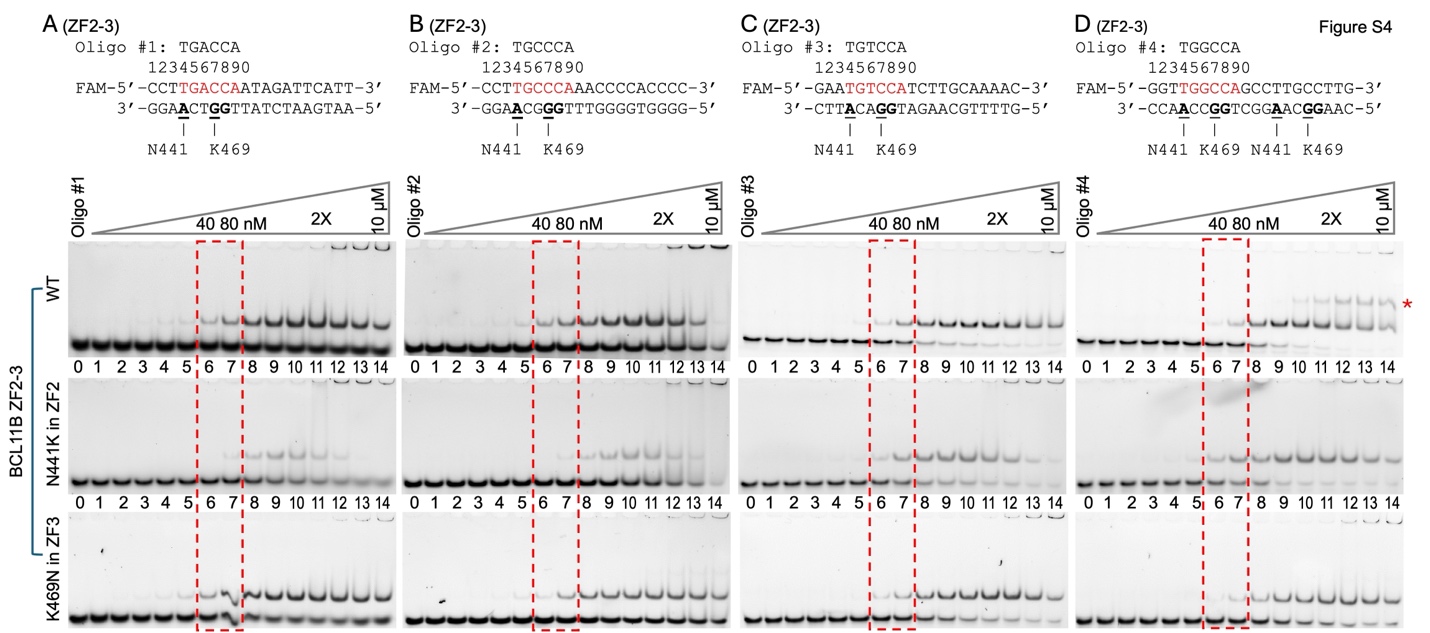

**Figure S4**, **EMSA analysis of** **N441K and K469N mutants in BCL11B ZF2-3.** (**A-D**) **EMSA** using four 20-bp oligonucleotides (each at 20 nM; sequences shown above) with three protein variants, WT ZF2-3 (top panels), N441K (middle panels), and K469N (bottom panels). Protein concentrations were prepared as twofold serial dilutions starting from 10 μM. At higher protein concentrations (lanes 12-14), protein precipitation is observed in the loading wells.

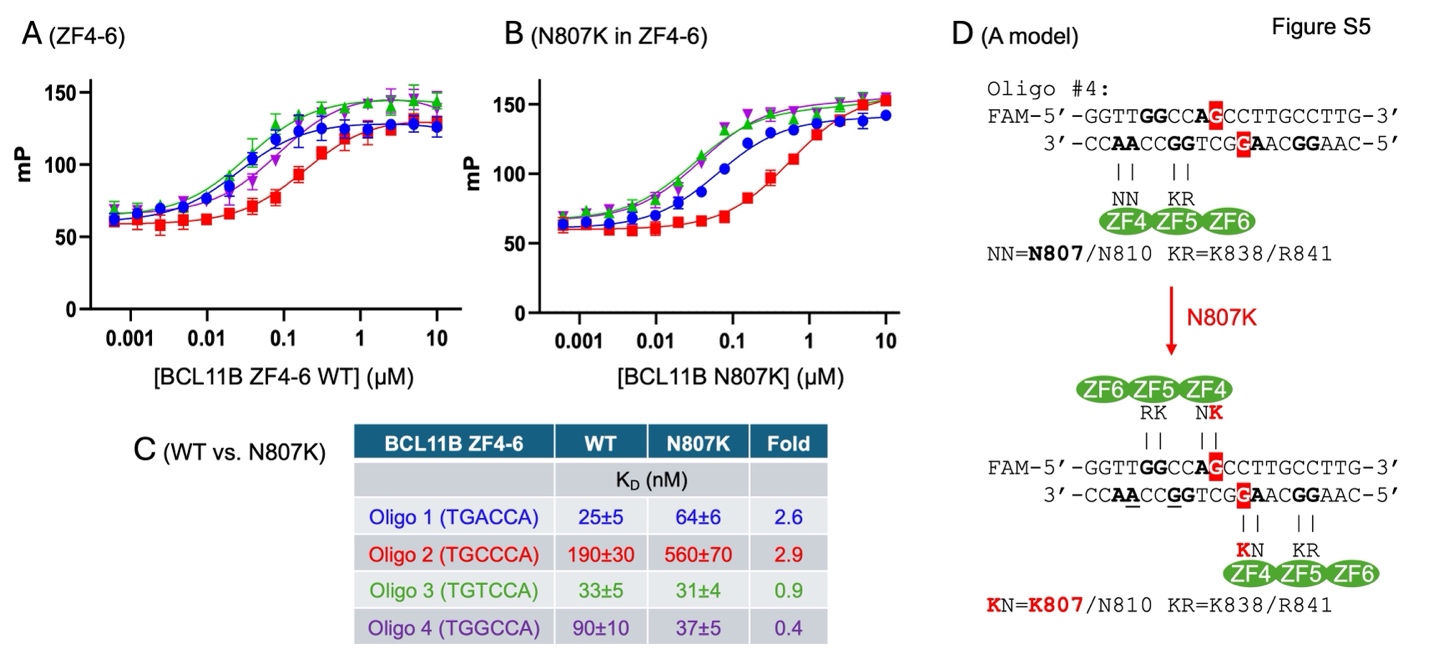

**Figure S5**, related to Figure 4, **FP analysis of WT (A) and N807K mutant (B) in BCL11B ZF4-6**. (**C**) Summary of K_D_ values and fold change between ZF4-6 WT and N807K mutant (Error bars represent SEM from N = 4 independent measurements). Colors shown here also apply to (A) and (B). (**D**) A model of N807K mutant changed specificity from N807-interacing Ade to K807-interacting Gua (highlighted in red). Two possible binding modes on either top strand or bottom strand are presented.

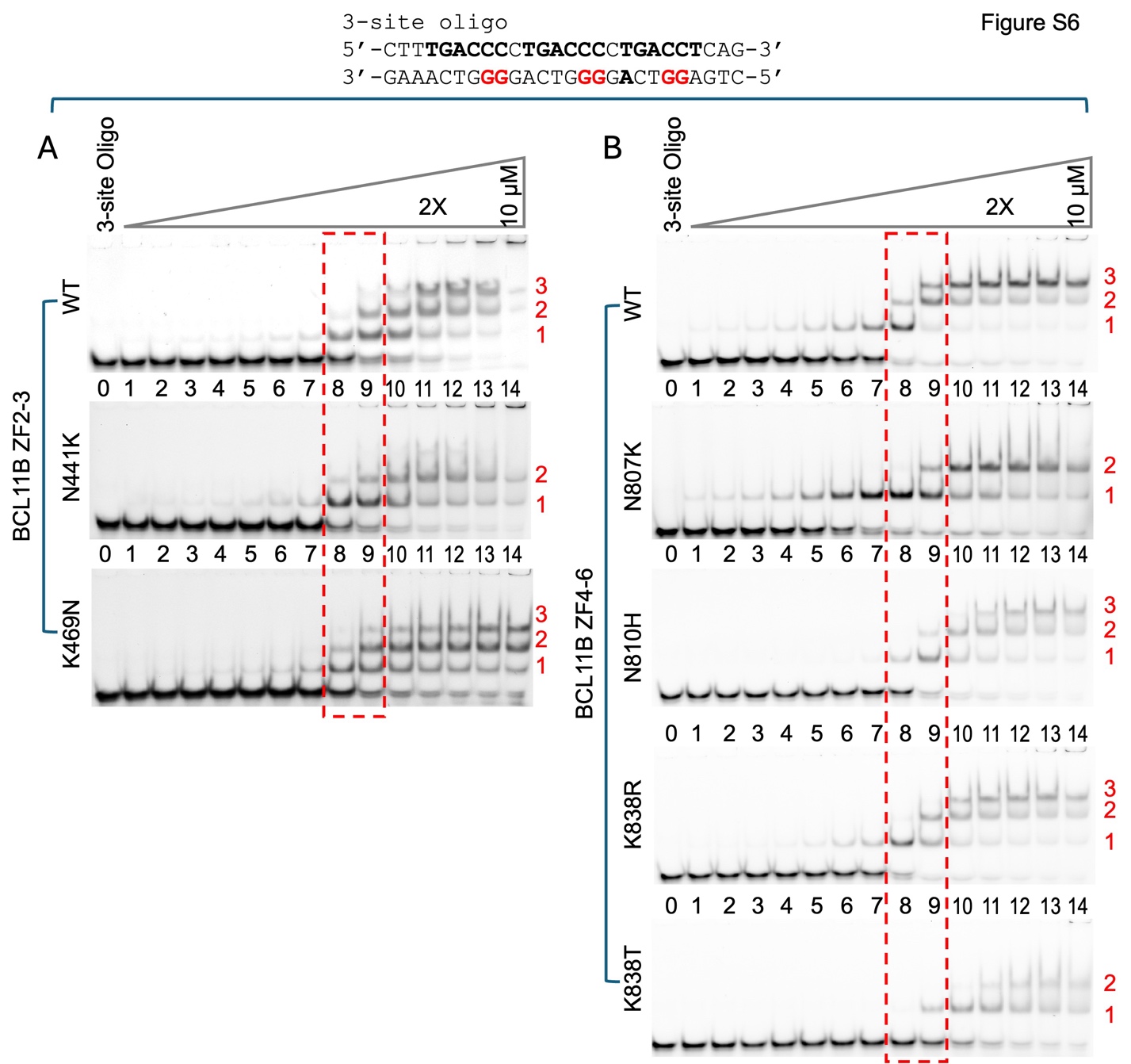

**Figure S6**, related to Figure 6, **BCK11B mutagenesis on 3-site oligo**. (**A**) EMSA of ZF2-3 WT, N441K and N469K. (**B**) EMSA of ZF4-6 WT, N807K, N810H, K838R, and K838T.

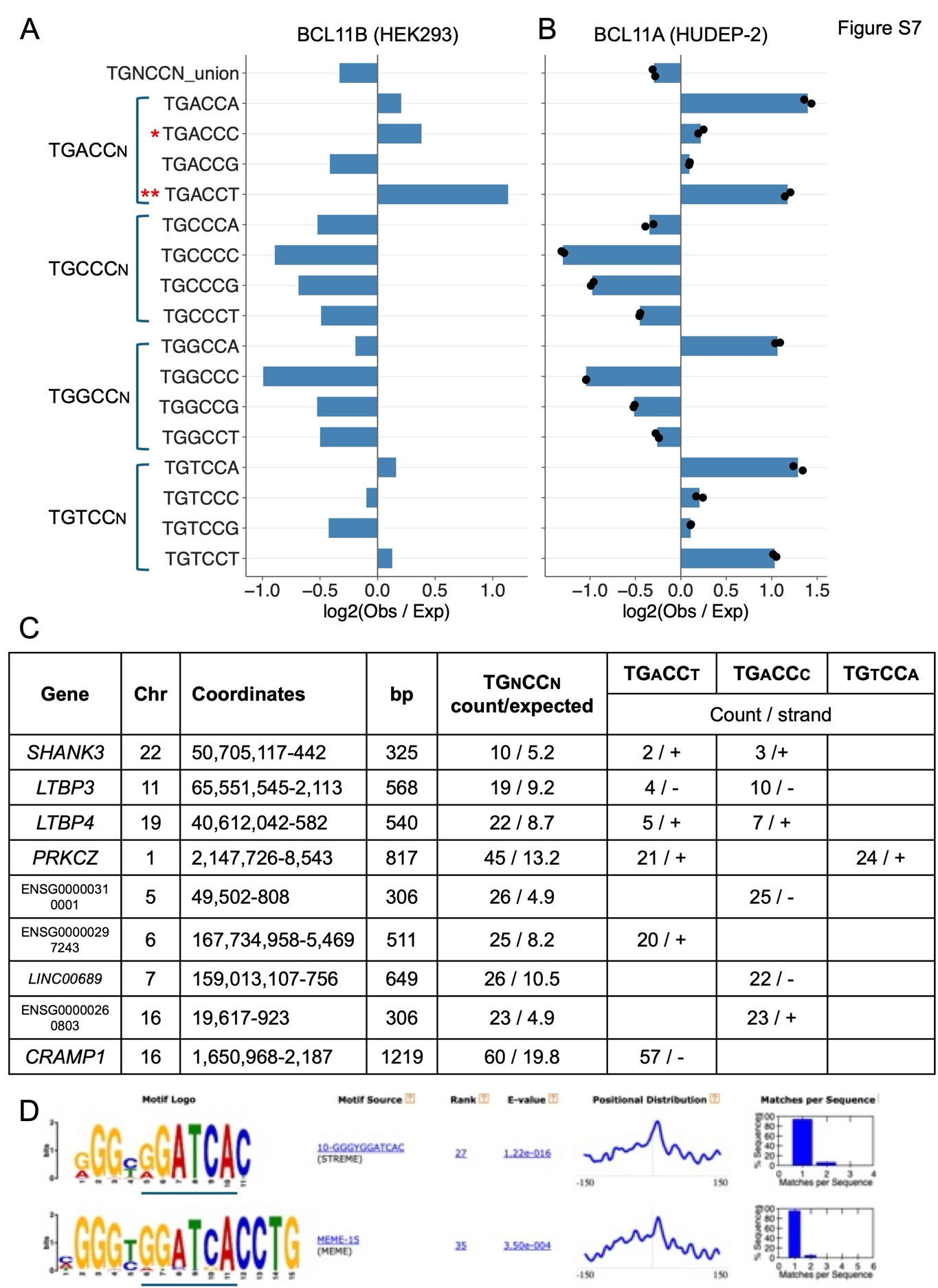

**Figure S7**, related to Figure 5H. Motif analysis of ENCODE ChIP-seq data for eGFP-BCL11B in human HEK293 cells (ENCSR770PQN). (**A**) Bar graphs show the log₂(observed/expected) enrichment of 6-mer TGnCCn motifs in eGFP-BCL11B ChIP-seq peaks, calculated relative to a first-order Markov background model derived from the same peak sequences. Values represent totals aggregated across all peaks. The union set (TGnCCn_union) combines counts across all 16 possibilities of TGnCCn, including reverse complements, with palindromic sequences counted only once. (**B**) For comparison, the same analysis was performed on ChIP-seq data for BCL11A-ER-V5 in HUDEP-2 cells ^4^. **(C)** TGnCCn motif enrichment in eGFP-BCL11B binding peaks associated with *SHANK3*, *LTBP3*, and *LTBP4*. TGnCCn motifs are enriched over expected based on Markov model. Specifically, more than half of the TGnCCn motifs in these peaks are TGACC(c/t) sequences and display strand specificity. This strand bias is both strong and consistently observed across many eGFP-BCL11B CHIP-seq peaks. In some regions, we observed extensive motif clustering, including 21 motif copies of TGaCCt and 24 copies of TGtCCa at the *PRKCZ* locus on chromosome 1 and up to 57 copies of TGaCCt at the *CRAMP1* locus on chromosome 16 within a single binding region, often in sequences enriched for repetitive elements. (**D**) De novo motif discovery also identified GGATCA motif, or its complimentary sequence, TGA**T**CC – a variation of TGa**C**Cc. This motif was also previously reported in ChIP-seq data obtained using an anti-Bcl11b antibody in transfected mouse striatal (STHdh) cells ^5^. In that study, Tang et al. ^5^ identified 1,410 significantly enriched Bcl11b-binding regions, of which only 36 were located within 1 kb of the TSS, indicating that most Bcl11b binding occurs distal to the proximal promoter regions.

**Table S1**. Summary of crystallization conditions for ZF2-3 and DNA complexes. Oil-like droplets were frequently observed in conditions that did not yield crystals (left panel).

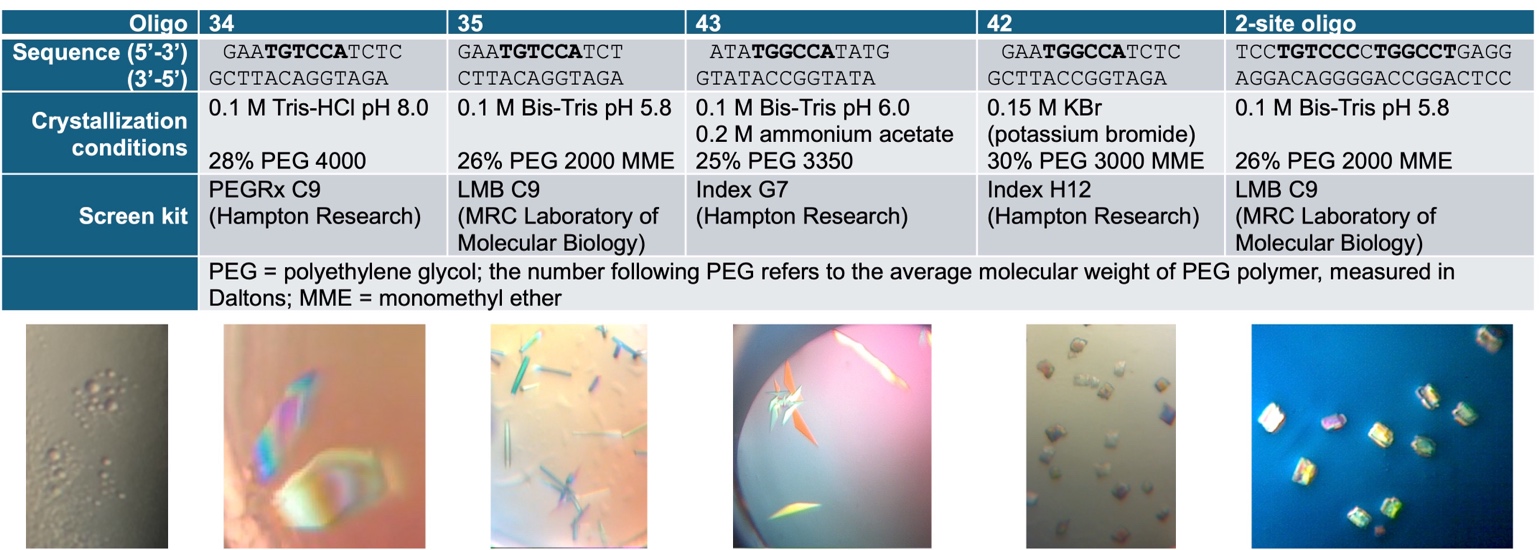

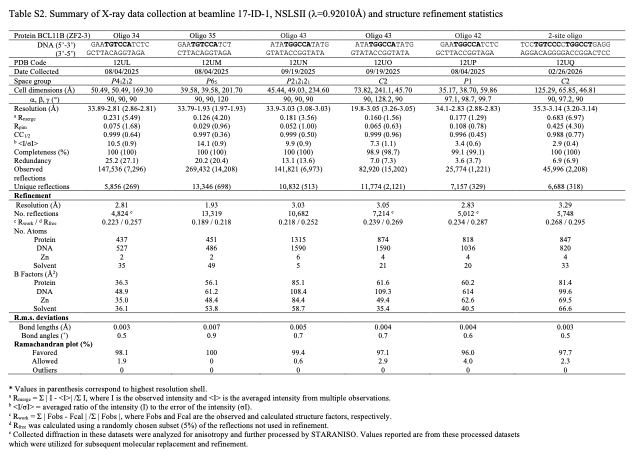
